# Genomic epidemiology of conflict-associated *Klebsiella pneumoniae* in Ukraine reveals a high-risk clone with a hybrid plasmid vulnerable to phages

**DOI:** 10.64898/2026.01.21.700813

**Authors:** Sabrina Green, Maha Niazi, Anatoliy Kazmirchuk, Olha Rubchuk, Olena Sanko, Viktor Tretiakov, Alessandra Carattoli, Pierre Bogaerts, Régis Tournebize, Valerie Lannoy, Catherine Eckert, Elisa D’Aliesio, Pieter-Jan Ceyssens, Diana Isabela Costescu Strachinaru, Griet Steurs, Christel Cochez, Johann Griselain, Maya Merabishvili, Anke Stoefs, Karlis Racenis, Volker Alt, Jeroen Wagemans, Thao Lyn Lê, Eveline Vriens, Rob Lavigne, Jean-Paul Pirnay

## Abstract

As antimicrobial resistance continues to rise and the development of new antimicrobials lags, we face an urgent threat where routine infections could become life-threatening. Armed conflicts, like Russia’s war on Ukraine, intensify this crisis by accelerating the emergence and spread of resistant organisms across regions and borders. Phage Support Ukraine (Phage Sp UKR) is a collaborative effort to provide phages targeting circulating pathogens in Ukraine for delivery of personalized phage therapies. Within this context, we received multidrug-resistant bacterial isolates belonging to a number of different species. We here focus on *Klebsiella pneumoniae*, the most concerning pathogen with regard to prevalence, virulence and antibiotic resistance. Among the *K. pneumoniae* isolates, we identified sequence types (ST) 39, 101, 147, 307 and 395, as well as a predominating clonal group—CG10146 (ST23 KL57), first detected in Moscow—carrying a hybrid virulence–resistance plasmid encoding NDM-1 and aerobactin. Over two years (2023–2025), this clonal group expanded and acquired pan-resistance. When tested in virulence assays, only one isolate (ST395) displayed hypervirulence in a *Galleria mellonella* though not in a mouse infection model. Several phages isolated from Ukrainian sewage were able to lyse these *K. pneumoniae* strains, including strains obtained from soldiers displaced in Belgium, Germany, and Latvia. Bacterial resistance was observed during *in vitro* testing. However, some phage-resistant isolates had mutations in virulence factors, including one that completely lost its hybrid plasmid, resulting in restored antibiotic susceptibility. Strategies like Phage Sp UKR are essential to prevent the selection, persistence, and global spread of these MDR clones.

## Introduction

Antimicrobial resistance (AMR) is the most significant threat to global health of our time. In the United States alone, 2.8 million antimicrobial-resistant infections occur annually^1^. In the European Union, AMR is responsible for 35,000 deaths each year, with additional healthcare costs totaling €1.5 billion^2^. A recent report estimated that bacterial AMR contributed to 1.27 million deaths worldwide in 2019, with highest burden in low- and middle-income countries (LMICS)^3^. The largest surge in AMR-related deaths was driven by pathogens resistant to carbapenems which totaled 1.16 million deaths in 2021^4^. The now updated 2024 prediction estimates that AMR could cause or contribute to 10 million deaths by 2050^4^.

Since the onset of the war on Ukraine, increased surveillance has detected rising AMR rates in European countries treating wounded soldiers. Phage therapy, or the use of bacterial viruses as a treatment for infections, has a storied history since its first use in 1919. In military medicine, phage therapy has previously been employed for soldiers and civilians suffering from bacterial infections prior to and after the development of antibiotics^5^. Today, extensive case studies and series indicate that phage therapy is safe and effective for difficult-to-treat infections when given in combination with antibiotics^6,7^.

Phage Support Ukraine (Phage Sp UKR) was born out of necessity, a collaboration between scientists, clinicians and military personnel to provide phages targeting circulating multidrug-resistant (MDR) strains in Ukraine-based hospitals. This initiative also supplies the essential equipment and training necessary to work with the phages and implement phage therapy for patients with difficult-to-treat infections. As part of this project, *Acinetobacter baumannii*, *Pseudomonas aeruginosa*, *Proteus mirabilis*, *Providencia stuartii*, and *Klebsiella pneumoniae* isolates were collected. Initial tests identified *K. pneumoniae* as the most concerning pathogen due to its antibiotic resistance, virulence, and prevalence among the isolates.

*K. pneumoniae* is a Gram-negative, encapsulated bacterium, linked to both hospital- and community-acquired infections, primarily affecting the immunocompromised. It can cause a wide range of infections, including pneumonias, urinary tract infections (UTIs), liver abscesses, bloodstream infections, surgical site and soft tissue infections, and sepsis. *K. pneumoniae* is the leading cause of neonatal sepsis in LMICs, attributing to 15% of all neonatal deaths^8,9^. The World Health Organization (WHO) has included carbapenem-resistant *K. pneumoniae* (CRKP) as a top bacterial priority pathogen as a way to address the pressing need for development of new, effective antibacterial therapies directed against this critical pathogen^3^.

*K. pneumoniae* is particularly adept at acquiring and disseminating antibiotic resistance, harboring multiple beta-lactamase genes often on transferable plasmids^10^. Some have described *K. pneumoniae* as having a “super-resistome”, conferring not only resistance to beta-lactams but also to aminoglycosides, quinolones, polymyxins and others^10^. These factors allow *K. pneumoniae* to thrive in healthcare settings, selected due to the continued widespread use of antibiotics. *K. pneumoniae* can be classified into two groups, classical *K. pneumoniae* (cKp) and hypervirulent *K. pneumoniae* (hvKp). The latter type is a major concern since hvKp strains are capable of causing severe, life-threatening infections in healthy individuals although treatable with antibiotics. This hypervirulent phenotype arises from a combination of virulence genes encoding for siderophores like aerobactin (*iuc*) and salmochelin (*iro*), as well as regulators of mucoid phenotype (*rmpADC* and *rmpA2*) which are found in the canonical hvKp virulence plasmid pLVPK ^11^. Recently there has been an emergence of new hybrid strains carrying both virulence and resistance genes, often within the same plasmid, with capsule types also associated with high morbidity ^12^. Distinguishing hvKp from cKp strains is critical to prevent the spread of MDR strains capable of causing severe disease.

A comprehensive assessment of the *K. pneumoniae* isolates was conducted to better understand the situation in Ukraine, focusing on their virulence and antibiotic resistance.

## Results

### Identification of *K. pneumoniae* isolates from Ukraine (UKR) as extensively drug-resistant (XDR) and multidrug-resistant (MDR)

The Queen Astrid Military Hospital (QAMH) in Brussels (Belgium) and the Laboratory of Gene Technology (LoGT) in Leuven (Belgium) received eight selected clinical isolates of *K. pneumoniae* directly from a military hospital in Kyiv, Ukraine. Susceptibility testing performed at QAMH, identified isolates UKR-KP3, UKR-KP4, UKR-KP5.1, UKR-KP5.2, and UKR-KP6 as extensively drug-resistant (XDR) defined as non-susceptible to at least one agent in all but two or fewer antimicrobial categories^13^. UKR-KP2, UKR-KP7, and UKR-KP8 as MDR. MDR was defined as non-susceptibility to at least one agent in three or more antimicrobial categories^13^ (Table 1).

**Table 1.**
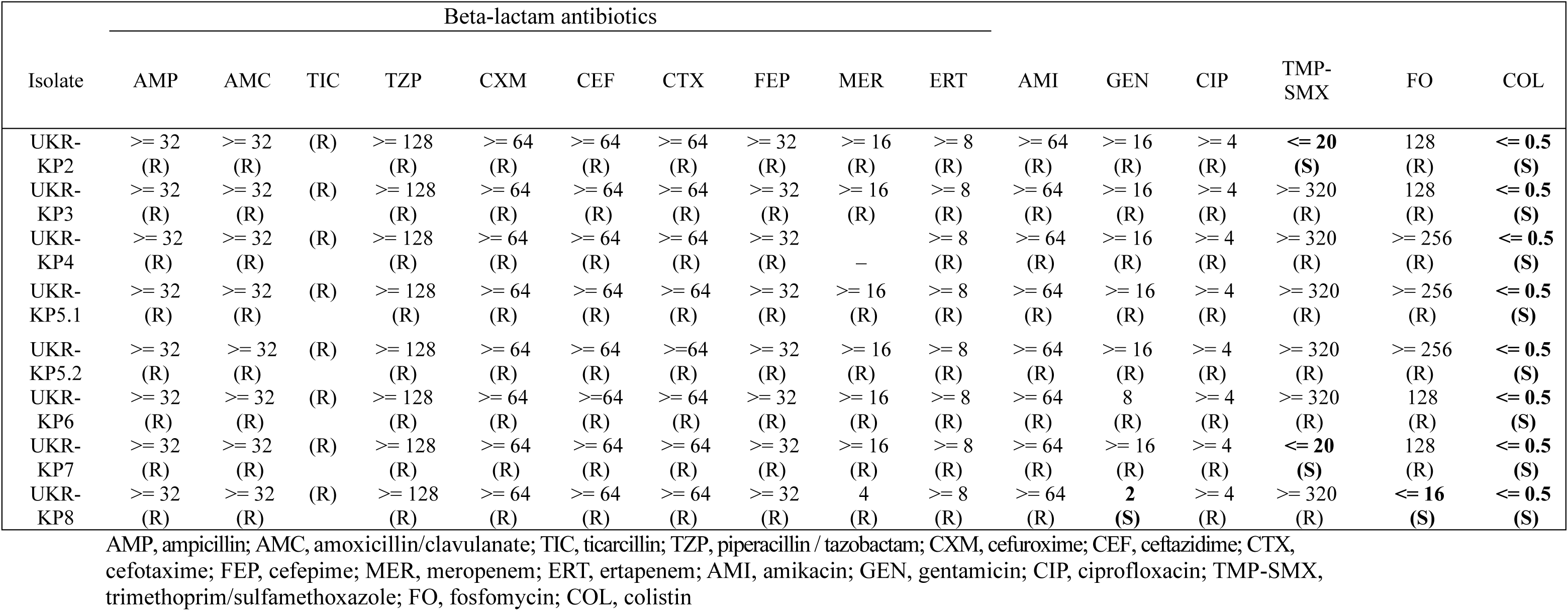
Antibiogram of UKR *K. pneumoniae* isolates.

All isolates tested showed non-susceptibility to the beta-lactam antibiotics tested, including penicillins (ampicillin and ticarcillin), penicillins with beta-lactamase inhibitors (amoxicillin/clavulanate and piperacillin/tazobactam), cephalosporins (cefuroxime, ceftazidime, cefotaxime, and cefepime) and carbapenems (meropenem and ertapenem). Moreover, all showed non-susceptibility to the fluoroquinolone ciprofloxacin. Most showed non-susceptibility to the aminoglycosides tested (amikacin 100% (8/8) and gentamicin 87% (7/8). Only UKR-KP8 was susceptible to gentamicin and fosfomycin. Two isolates, UKR-KP2 and UKR-KP7, were susceptible to the antifolates trimethoprim/sulfamethoxazole. All isolates were susceptible to the last-line antibiotic colistin.

Whole genome sequencing analysis showed several resistance genes to major antibiotic classes in agreement with the antibiotic susceptibility data (Supplementary Tables 1 and 2). All isolates contained at least three different beta-lactamase genes on plasmids (Supplementary Table 2). Some of these isolates harbored *bla*_KPC-2_, (UKR-KP2 and UKR-KP7), a class A serine beta-lactamase which confers resistance to penicillin, cephalosporins, and carbapenems. Other carbapenemases included *bla*_NDM-1_ (87%, 7/8) and *bla*_OXA-48_ (62%; 5/8) which have also been detected in other surveillance studies associated with the war on Ukraine^14,15^. For UKR-KP2 and UKR-KP7 we identified seven different beta-lactamases (*bla*_KPC-2_, *bla*_NDM-1_, *bla*_TEM-1B_, *bla*_CMY-4_, *bla*_CTX-M-15_, *bla*_OXA-1_, and *bla*_OXA-10_).

### UKR *K. pneumoniae* isolates from circulating MDR sequence types

Multilocus sequencing type (MLST) analysis assigned the isolates to sequence types (ST)23 (UKR-KP3, UKR-KP5.1, and UKR-KP5.2), ST39 (UKR-KP2 and UKR-KP7), ST307 (UKR-KP8), and ST395 (UKR-KP4 and UKR-KP6) (Fig. 1a). Multiple surveillance studies have similarly identified these sequence types in patients associated with the war in Ukraine^14–17^. The isolates were assigned the following K loci (KL): KL2, KL23, KL57, KL102, and KL108 (Supplementary Table1). The isolates were assigned a virulence score ranging from 1 to 5, based on the presence and combination of virulence loci (*ybt* = yersiniabactin, *iro* = salmochelin, *cbl* = colibactin, *iuc* = aerobactin^18^) (Supplementary Table 3). An additional genome analysis was performed using the AntiSMASH software to estimate the biosynthetic potential for secondary metabolites with results are outlined in Supplementary Table 4.

**Fig. 1.**
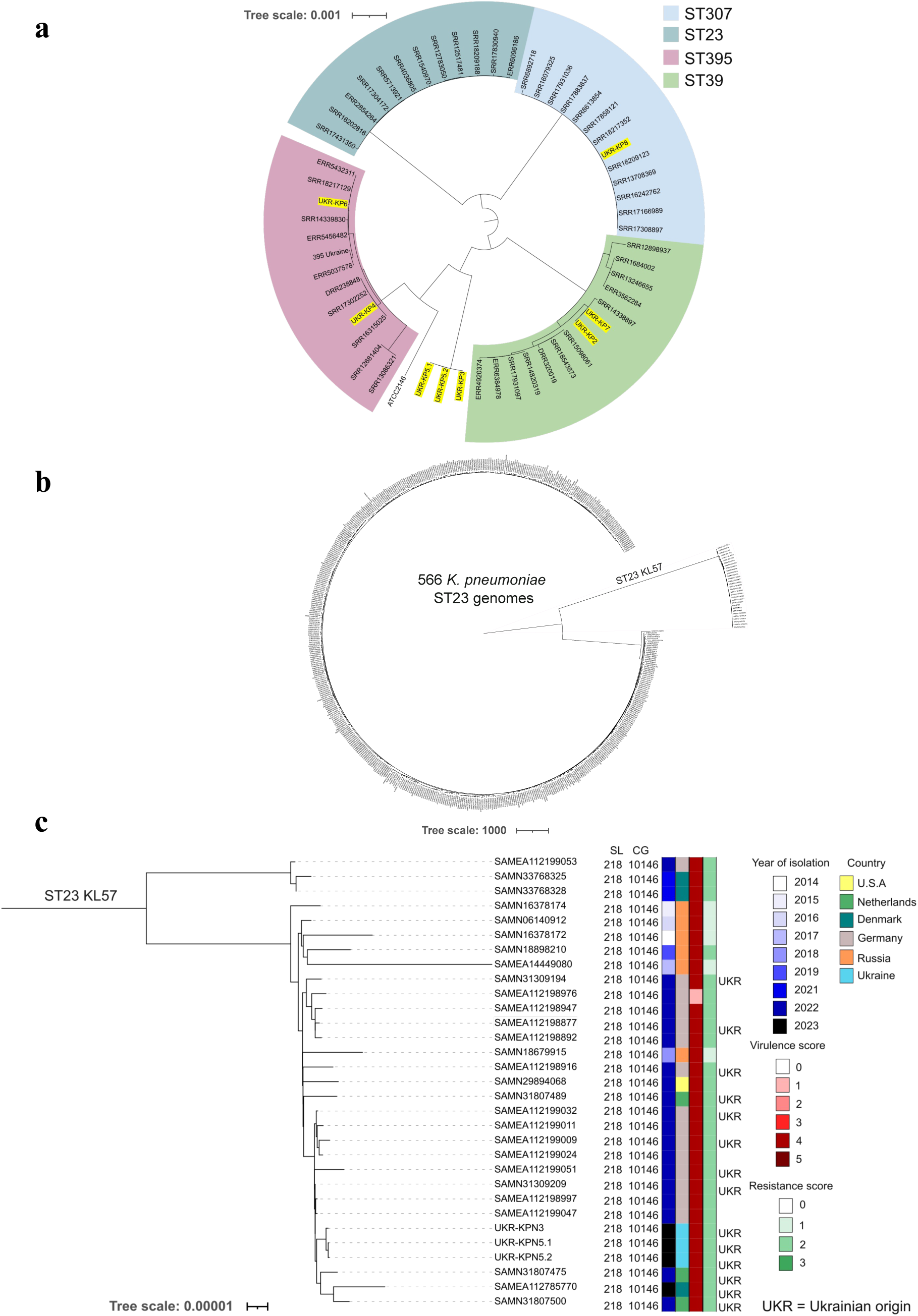
Phylogenetic analysis of *K. pneumoniae* isolates. **a**, Maximum likelihood phylogenetic tree of the genomes of the analyzed sequential bacterial isolates based on a core gene alignment. Highlighted in yellow are the Ukrainian isolates. **b**, All ST23 genomes present in Pathogenwatch (accessed July 2024) including the Ukrainian isolates. **c**, A phylogenetic tree of the ST23 KL57 genomes that deviated from the ST23 group. Year of isolation, country, virulence and resistance score were determined using Pathogenwatch which uses the Kleborate tool. The defined SL (sublineage) and CG (clonal group) were determined using the *Klebsiella* Pasteur institute MLST database. UKR (Ukrainian) origin was confirmed by a literature search by accession # provided by Pathogenwatch.

Overall, biosynthetic gene clusters (BGC) encoding siderophores and encoding ribosomally synthesized and post-translationally modified proteins (RiPPs) associated with virulence and interspecies competition were detected in all isolates.

The Kleborate genome screening tool also detected the presence of capsule up-regulation genes *rmpADC* and *rmpA2*. For two isolates, UKR-KP7 and UKR-KP2, these loci were not detected. For the others, if this was detected, a frameshift mutation in *rmpA*/*rmpA2* was noted leading to a truncation in the protein.

A phenotypic evaluation determined none were hypermucoviscous UKR-KP3, UKR-KP5.1, and UKR-KP5.2 produced a 5 mm string, which is below the typical cut-off for positive string test (data not shown).

A dendrogram representing the sequence types ST23, ST39, ST307, and ST395 using *K. pneumoniae* strains from the website Pathogenwatch showed that isolates UKR-KP3, UKR-KP5.1, and UKR-KP5.2 that were initially determined to be ST23 did not cluster with other representatives from this sequence type (Fig. 1a). All other strains did cluster within their initially assigned ST groups. Further analysis using all ST23 *K. pneumoniae* genomes from Pathogenwatch (566 genomes) also showed that our three ST23 isolates clustered away from this sequence type into the ST23-KL57 cluster alongside 28 other isolates (Fig. 1b). Among these 31 ST23-KL57 isolates (Fig. 1c) metadata indicated that a majority (23/31) were recently isolated (2022–2023). Most had a high virulence score of 4 (30/31) and a high antimicrobial resistance score of 2 (out of 3) (26/31). Interestingly, almost half of them (14/31) have some link with Ukraine. These isolates resulted from surveillance studies of displaced Ukrainians in Germany (SAMN31309194, SAMEA112198877, SAMEA112198916, SAMEA112199032, SAMEA112199009, SAMEA112199051, and SAMN31309209), the Netherlands (SAMN31807489, SAMN31807475, and SAMN31807500), and Denmark (SAMEA112785770)^14,15,17,19^. Also, six out of 17 isolates for which no Ukrainian origin could be documented were isolated in Russia. These isolates of Russian origin were isolated at earlier time points (2014–2019) and had a high virulence score of 4 (6/6) but 5/6 had a lower resistance score of 1.

ST23-KL57 isolates have been reported in other studies regarding the AMR problem as well^12,20^. As expressed in other studies, the 7-locus MLST we employed is not sufficiently discriminatory to distinguish clonally related *K. pneumoniae*^21^. A core-genome (cg) MLST analysis using the BIGSdb-Pasteur database showed that all ST23-KL57 isolates (UKR-KP3, UKR-KP5.1, and UKR-KP5) as well as the 28 others that clustered away from the ST23 genomes belong to the sublineage (SL)218 clonal group (CG)10146.

### Plasmid content of UKR isolates associated with virulence and antibiotic resistance

Within the genomes of the CG10146 group we identified eight plasmids (Supplementary Table 5). This group carried a large (243 kb in UKR-KP5.1) plasmid derived from the hvKp virulence plasmid pLVPK. Figure 2a and b shows a graphical comparison of the plasmids identified in isolates UKR-KP3, UKR-KP5.1, and 5.2 (not shown) compared to the canonical hvKp plasmid pLVPK^22^. The UKR isolates carry an additional *bla*_NDM-1_ not present in pLVPK on their hybrid plasmid while maintaining the replicon (Rep) VIR, the tellurite and silver and copper resistance clusters, and four clusters involved in iron and other ions transportation (including aerobactin), which are usually present on pLVPK. Some virulence genes are attenuated by several mutations including a frameshift in *rmpA2* regulator gene, and the *rmpAD* gene and the *iro* and *fec* clusters are lacking. There is also a change of one replicon. Plasmid pLVPK usually harbours both Rep_VIR (typical of the virulence plasmid) and the HIB (pNDM-Mar) replicon, identified for the first time in a plasmid called pNDM-Mar. Instead, this pLVPK-like plasmid carries the Rep_VIR but not the HIB (pNDM-Mar) replicon, which is substituted by a pC39 replicon that was not previously present in PlasmidFinder. This replicon derives from another type of plasmid identified in China as pC39_CP061702.1. The difference between the pLVPK-like plasmids of UKR-KP3, UKR-KP5.1, and UKR-KP 5.2 is the presence of an ArmA (aminoglycoside resistance methylase) region, inserted in the resistance region carrying *bla*_NDM-1_.

**Fig. 2.**
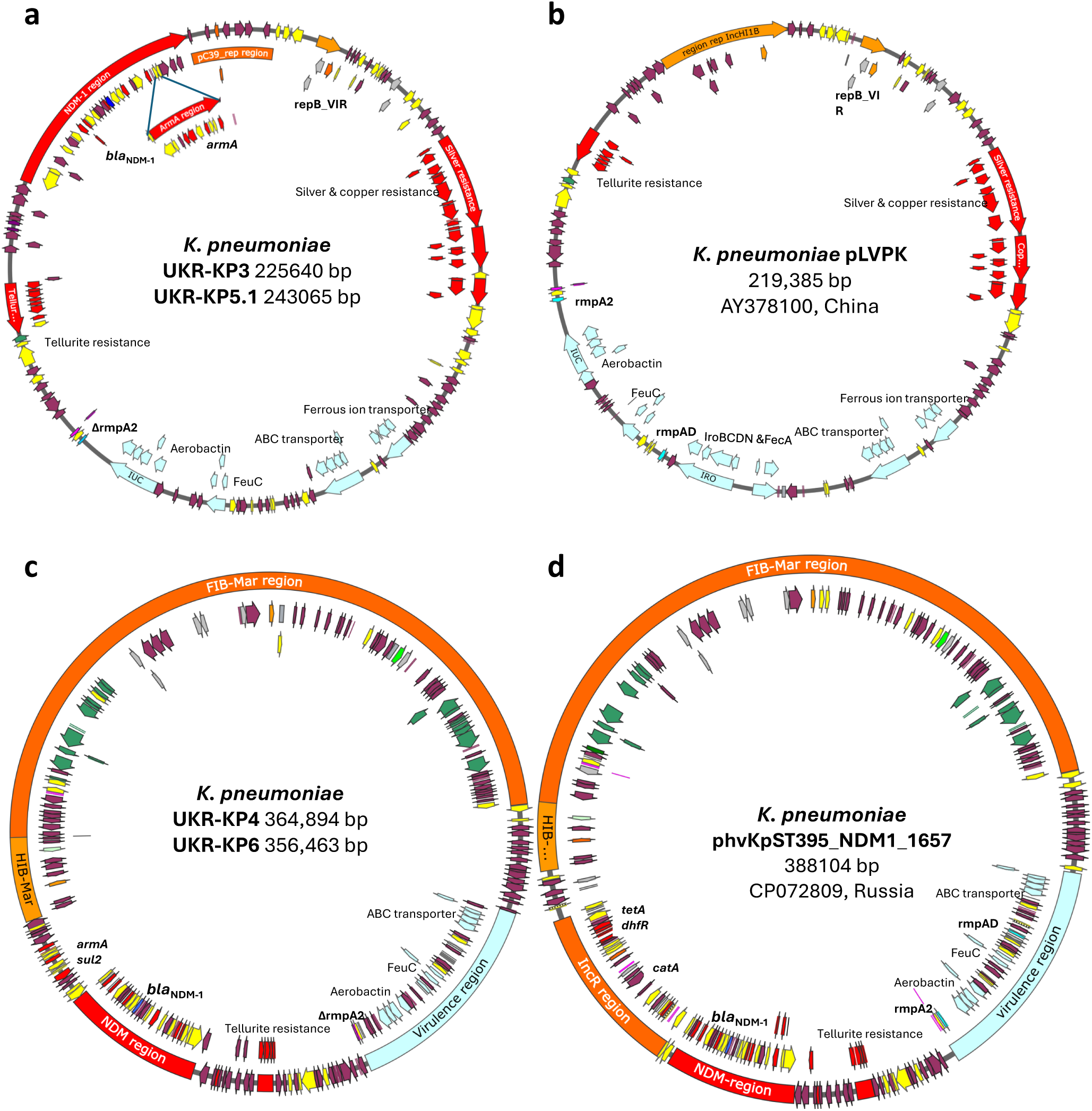
Virulence plasmids of *K. pneumoniae* ST23 and ST395. a, Plasmid present in UKR-KP3, KP5.1 in comparison with **b,** the hypervirulence plasmid pLVPK (accession # AY378100) described in China in 2004. UKR-KP3.1 is depicted in the figure, in plasmid UKR-KP5 there is an additional *armA* region. **c**, Plasmid present in ST395 isolates (UKR-KP4 and UKR-KP6) in comparison with (**d**) plasmid phvKpST395_NDM1_1657 (accession # CP072809) identified in *K. pneumoniae* ST395 in Russia in 2021. The major features are highlighted in: red, resistance genes; pale blue, virulence regions; orange, replicon regions; green, transfer loci for conjugation; yellow, mobile genetic elements, insertion sequences, integrase of class 1 integrons; violet, hypothetical proteins. *rmpA2* and *rmpA,B,C,D* gene positions are in bold.

For the ST395 isolates, UKR-KP4 had three plasmids and UKR-KP6 had four. Interestingly, both carried a hybrid resistance and virulence plasmid highly related to a plasmid described in Russia (phvKpST395_NDM1_1657, Fig. 2c and d)^23^. The unique difference is the lack of the IncR region where there is an acquired *armA* gene. This plasmid carries the HIB replicon (the same of pLVPK) associated in the original pNDM-Mar plasmid with a FIB(Mar) replicon also carrying two conjugation loci. These large plasmids carry several virulence factors including aerobactin and other iron transporters^23^.

The ST39 isolates UKR-KP2 and UKR-KP7 both carried six plasmids. A new small plasmid was identified in both. This plasmid, named ColpB8919, was found previously in strain SCPM-O-B-8923 isolated from patients in Moscow, Russia. The UKR-KP8 isolate had three plasmids outlined in Supplementary Table 5.

### Virulence phenotype of UKR isolates

Given the presence of virulence genes in these isolates, we assessed their pathogenicity using the wax moth (*Galleria mellonella*) larvae model, which has been widely used to study *K. pneumoniae* virulence^24,25^. A challenge of (1–5) X 10^4^ CFU was used as this dose causes the rapid death (within 48 hours) of larvae by virulent strains of *K. pneumoniae* in this model system^24,26^. Within six hours post-infection the UKR-KP6 infected larvae group exhibited low activity scores (Fig. 3a). This dose led to rapid and acute death in this group (79% mortality within 24 hours) (Fig. 3b). Some deaths were noted in the other groups as well, including the UKR-KP2 and UKR-KP7 groups (∼32% in 24 hours) and the UKR-KP4 and UKR-KP8 groups (∼15% mortality in 24 hours). No deaths were noted in the groups infected with UKR-KP3, UKR-KP5.1, and in the PBS control group. UKR-KP5.2 was not included in this assay as this bacterial isolate was genomically identical to UKR-KP5.1.

**Fig. 3.**
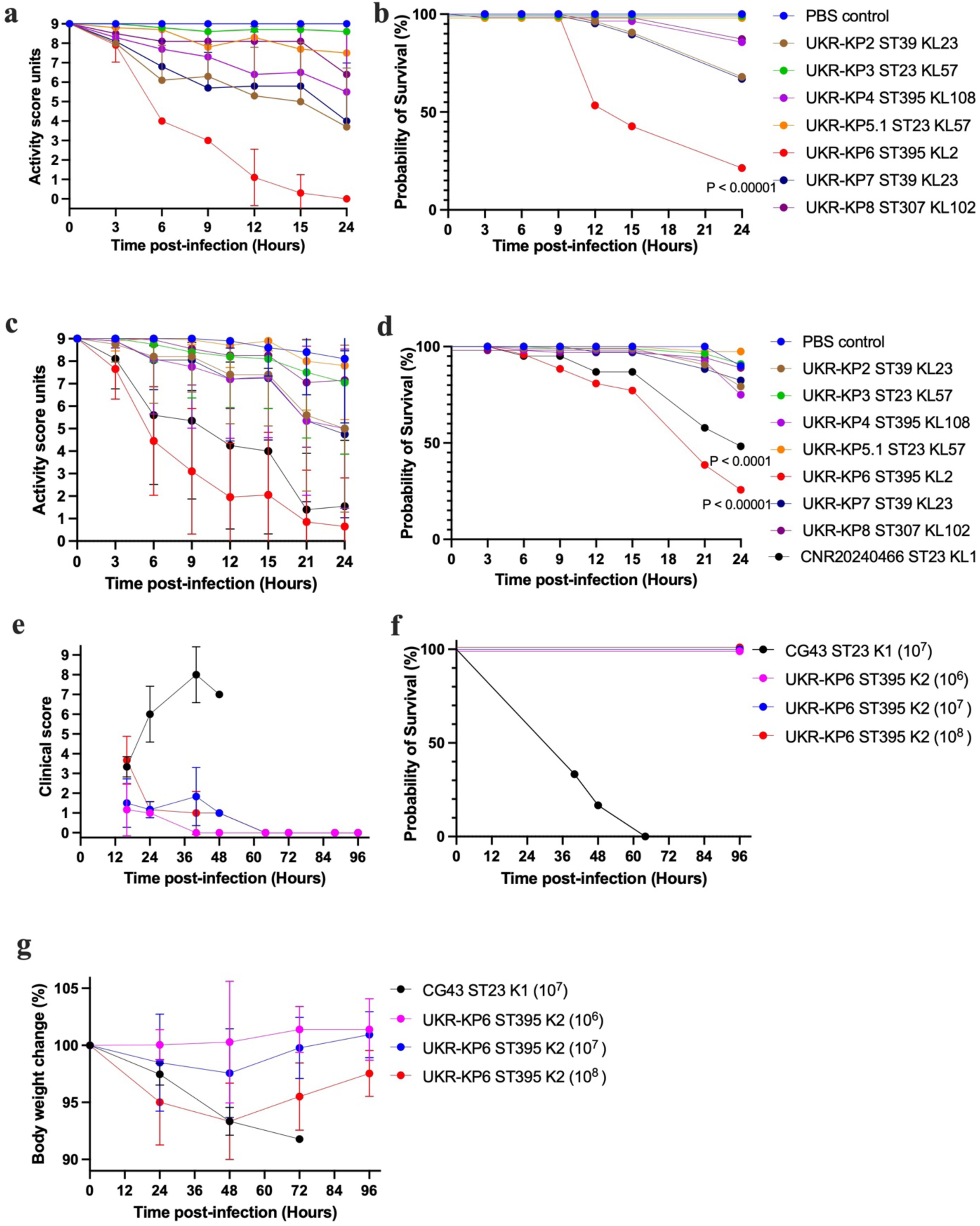
Virulence assessment of Ukrainian *K. pneumoniae* isolates. a, *Galleria mellonella* larvae were injected with *K. pneumoniae* isolates at a dose of 10^4^ CFU or with PBS and incubated at a temperature of 37°C. The larvae were monitored every 3 hours to assess for activity. **b,** Probability of survival **c,** *G. melonella* were similarly infected and assessed for activity with the inclusion of a hypervirulent *K. pneumoniae* strain CNR20240466 as a control. **d,** Probability of survival. **e,** BALB/cJRj female and male mice were anesthetized and infected intranasally with either 10^6^, 10^7^, and 10^8^ CFU of UKR-KP6. The control CG43 was infected with 10^7^ CFU. Activity was assessed twice daily. **f,** Probability of survival. **g,** Weight assessment of mice. Values (dotes) Clinical score was assessed twice daily. Values (dotes) indicate the mean. For the Galleria experiments N=10. For the mouse experiments N=6 (3 males and 3 females). P values were determined using the log-rank test using Prism 10

This experiment was repeated twice with the inclusion of a strain with the highest virulence score of 5 with the intact *rmpADC* but not *rmpA2*-ST23 KL1 strain CNR20240466. Around six hours post-infection, a precipitous drop in activity scores was observed for the group infected with UKR-KP6 as well as the group infected with the ST23 KL1 strain (Fig. 3c). At 24 hours post-infection more deaths were noted in the UKR-KP6 (∼75% mortality) compared to the ST23 KL1 strain (∼52% mortality) (Fig. 3d). Both groups showed statistically significant lower values than the PBS control. Of note, one *G. mellonella* larva died in the PBS control group likely due to a technical error. All other groups clustered together and did not show a statically significant difference from the control.

To further evaluate the virulence of this UKR-KP6 isolate we utilized a mouse model of pneumonia. BALB/cJrj male and female mice were anesthetized and infected intranasally at different dosages with clinical scores assessed twice daily and mice weighed daily to assess health. Despite its high virulence in *G. mellonella*, UKR-KP6 was avirulent in the mouse model when infected with 20 µL of 10^6^, 10^7^, and 10^8^ CFU (Fig. 3 e-g). In contrast, another ST23 KL1 strain (with a virulence score of 4) was used as a control for this model system causing 100% mortality within 60 hours at a 10^7^ dose (Fig. 3f). The disease activity index and body weight of UKR-KP6-infected mice remained stable, with consistently low disease scores throughout the infection (Fig. 3 e-g).

### Further expansion of CG10146 in neighboring countries and updated UKR 2025 isolates

A concomitant surveillance study was performed on phenotypic and genotypic characteristics of bacteria isolated from Ukrainian war victims treated in the Burn Wound Center of the QAMH (Strachinaru et al., *unpublished*). We received additional isolates from UKR soldiers displaced in Latvia and Germany. A phylogenetic tree showed that many of the isolates from these soldiers grouped in with the isolates we received directly from Ukraine (Fig. 4. These included the sequence types ST395 (four isolates) and CG10146 (four isolates).

**Fig. 4.**
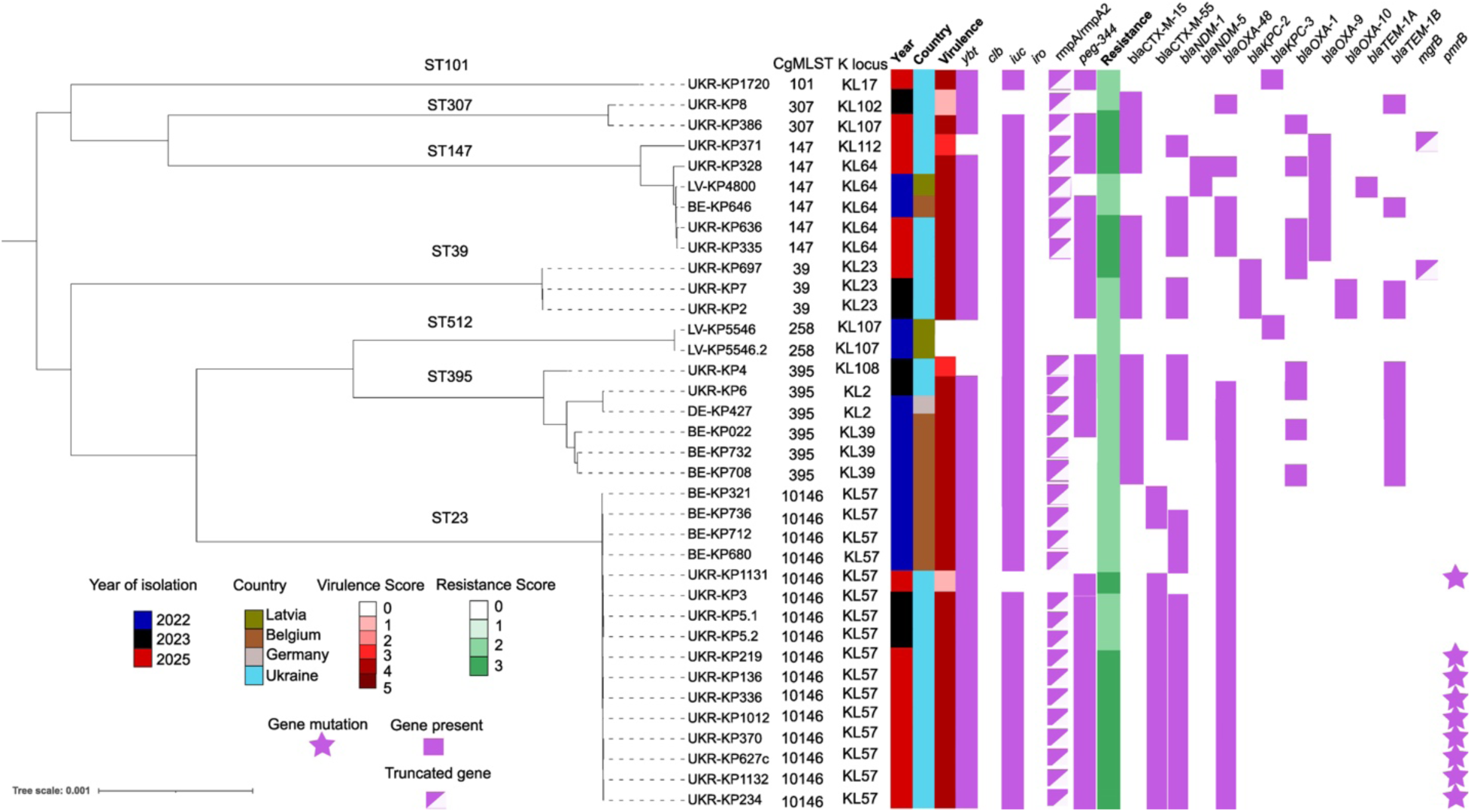
Phylogenetic analysis of *K. pneumoniae* isolates from injured Ukrainian soldiers in Belgium and in Ukraine and isolated from Kyiv in 2025. A maximum likelihood phylogenetic tree of the genomes of the analyzed sequential bacterial isolates based on a core gene alignment. Resistance and virulence genes were determined using Pathogenwatch. *ybt*= Yersiniabactin, *clb*= Colibactin, *iuc*= Aerobactin, *iro*= Salmochelin, *rmpArmpA2* = regulator of mucoid phenotype, *bla* = beta-lactamase gene.

In 2025, 16 additional isolates were received, including 13 CG10146 isolates, bringing CG10146 to 16 isolates in total. A notable difference in this expanded set was that all CG10146 isolates had acquired resistance to colistin (Table 2) and still carried multiple beta-lactamases and antibiotic resistance genes carried in plasmids (Fig. 4 and Supplementary Tables 6 and 7). Although the *mcr-1* gene was not detected, mutations were found in *pmrB,* a gene associated with chromosomal colistin resistance. When comparing to the others in this clonal group, specifically, a single SNP resulting in a Thr157Pro substitution was identified, a mutation previously shown to be sufficient to confer colistin resistance^27^. In contrast, colistin resistance in isolates from other sequence types appeared to result from truncations in *mgrB*, another well-known resistance mechanism. However, we could not definitively determine the genomic mutation that led to colistin resistance in the others.

**Table 2.**
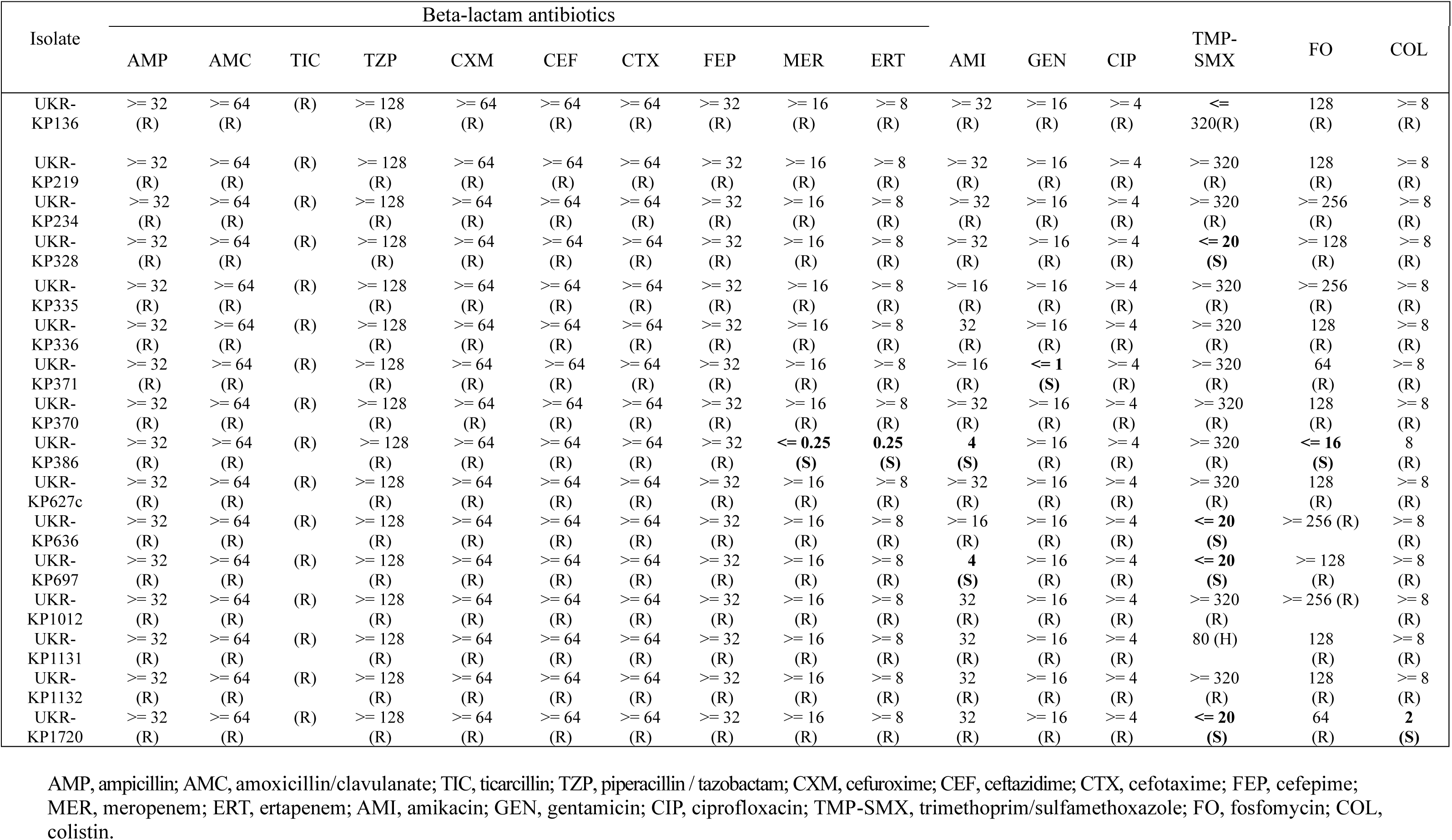
Antibiogram of UKR *K. pneumoniae* isolates 2025.

All isolates in CG10146 retained a high virulence score (3), except UKR-KP1131, which lacked NDM-1. These isolates also exhibited the highest resistance scores (3), reflecting carriage of multiple beta-lactamase genes in addition to colistin resistance. Notably, the *peg-344* gene—encoding a metabolite transporter associated with hypervirulence—was present in all newly assessed genomes in this clonal group^28^.

### Phage susceptibility of UKR isolates

The large QAMH phage collection was tested against these strains. However, only four phages (Fig. 5a; QAMH API’s) were able to infect a subset of them (7/37, 18%). Consequently, additional phages were isolated from Ukrainian wastewater, using UKR strains as hosts. A diverse panel of phages were isolated (n=31) and tested against all Ukrainian derived isolates (Supplementary Figure 1). The efficiency of plating (EOP) assay showed that all isolates could be infected by at least one phage. One phage was identified as temperate and was therefore excluded from further study. The remaining Ukrainian phages belonged predominately to the genus *Drulisvirus* (23/30), with additional representatives from *Efbeekayvirus* (2/30), *Webervirus* (2/30), *Slopekvirus* (1/30), *Pseudotevenvirus* (1/30) and *Przondovirus* (1/30).

**Fig. 5.**
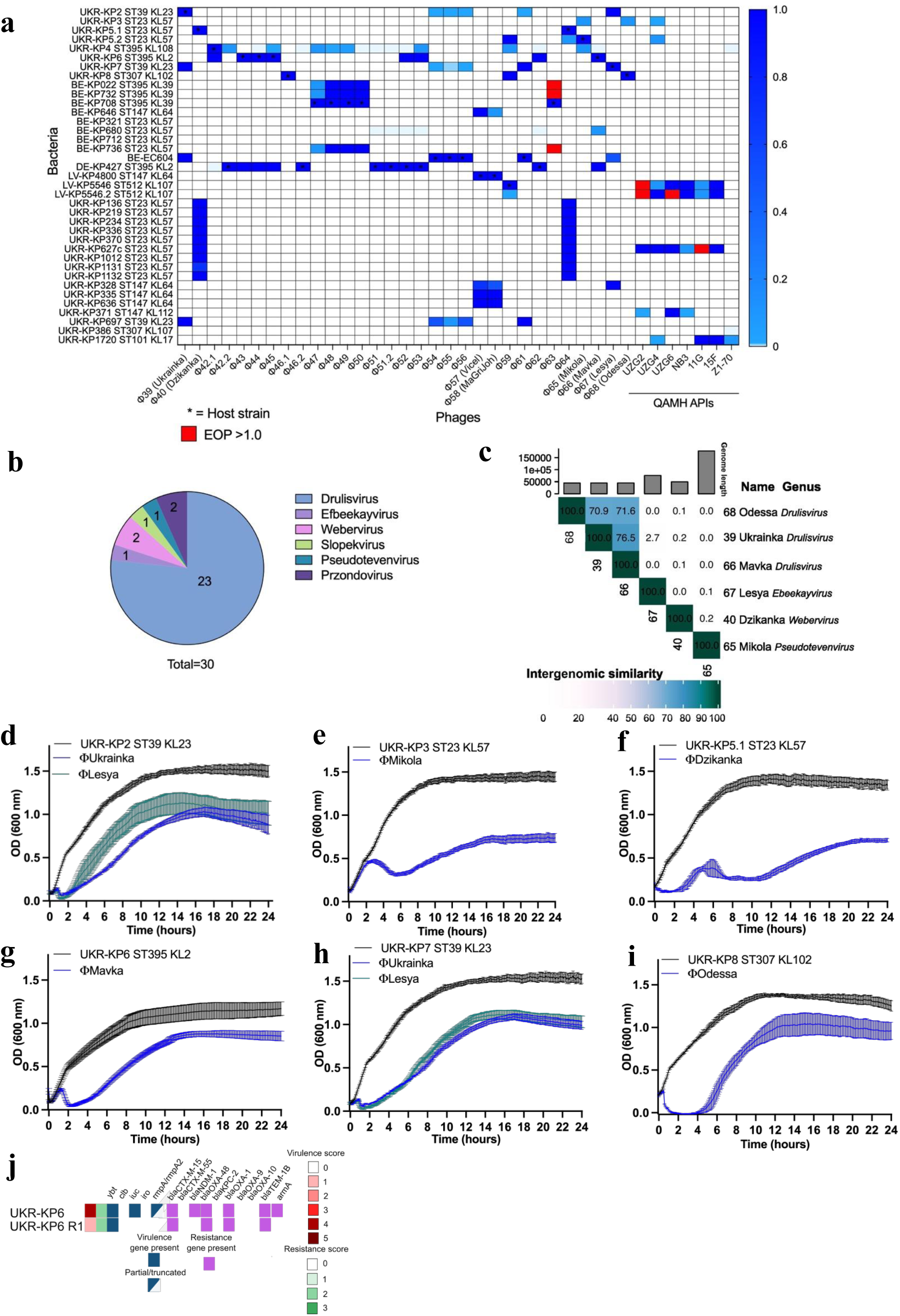
Phage assessment on Ukrainian isolates. a, Efficiency of plating (EOP) of phages on bacterial isolates from soldiers in Ukraine (UKR), Belgium (BE), Germany (DE) and Latvia (LV). Darker blue indicates higher EOP. *=host strain. Red = EOP > 1. **b,** abundance of different genera present in 30 UKR phages **c,** Intergenomic similarity of the UKR phages is represented in a VIRIDIC generated heatmap.**d-i,** Killing curves performed at a multiplicity of infection (MOI) of 1. Optical density readings taken every 5 minutes for 24 hours. Mean and standard deviation are represented of 3 biological replicates. **j,** Resistance and virulence change in a UKR-KP6 phage resistor. Genes were determined using Pathogenwatch.

The lytic activity of the selected phages against Ukrainian isolates representing major ST groups including the ST23-KL57 lineage, was further assessed in a 24-hour liquid culture assay. Six phages were the most effective at suppressing bacterial regrowth (Φ39, 40, 65, 66, 67, and 68) for 2 to 4 hours; however, regrowth occurred in all cases (Fig. 5c-i). The phages were renamed as Ukrainka, Dzikanka, Mikola, Mavka, Lesya, and Odessa. Further genomic similarity analyses are shown in Supplementary Table 8. All phages exhibited synergistic interactions with meropenem, resulting in the absence of bacterial regrowth for 48 hours, except for phages Lesya and Mikola (Supplementary Figure 2; data not shown).

### Resistance assessment of UKR phages

We isolated and sequenced phage-resistant bacteria after a 24-hour killing curve assay to identify mutations associated to phage resistance which could lead to fitness costs or could be linked to identifying the receptor of the phage. These mutations are listed in Supplementary Tables 9 and 10.

For the UKR-KP5.1 and UKR-KP8 resistors (resistant to phages Dzikanka and Odessa) mutations were only identified in hypothetical genes or in non-coding regions. In a UKR-KP2 resistor to phage Ukrainka, a SNP mutation was identified in *arnC*, encoding an undecaprenyl transferase involved in the lipid A modification pathway^29^. This mutation introduces a stop codon, leading to premature truncation of the protein. In a UKR-KP7 resistor to phage Ukrainka, one large deletion was identified in a region with genes important for capsule polymerization (*wzC*, w*zA*, w*zI*)^30^. In UKR-KP7 (resistant to Ukrainka) a deletion in a region with genes important for O-antigen biosynthesis (*ugd*, *rfbC*, *rfbD*, *rffH*, *rffG*, *gnd)*^30^ was identified. In a UKR-KP3 resistor to phage Mikola, a frameshift variant in the bacterial maltose pore LamB, receptor to phage λ, was identified^31^. Finally, in a UKR-KP6 resistor to phage Mavka, we identified a SNP mutation in the cps export gene *wzC*, leading to a missense mutation. In addition, a mutation was detected in another isolate within the capsule biosynthesis gene *wcaJ*. Interestingly, in some of these isolates, the large hybrid virulence and resistance plasmid (phvKpST395-like; Figure 3c) was lost. However, despite this isolate now having a virulence score of 1 instead of 4 (Figure 5j; yersiniabactin only present), no difference in virulence was noted, when tested in the *G. mellonella* model, compared to UKR-KP6 (Supplementary Fig. 2a and b). An antibiogram was performed on this isolate in which we now observed a re-sensitivity to the aminoglycosides amikacin and gentamicin (Table 3). We repeated the isolation of resistors on UKR6 and saw this phenotype again in total 3/9 (33.3% of the time) resistors in total (loss of plasmid). A plasmid stability assay showed this plasmid to be stable (96% retention in 2 replicates) suggesting loss of the plasmid may be a result of infection or resistance to the phage (data not shown).

**Table 3.**
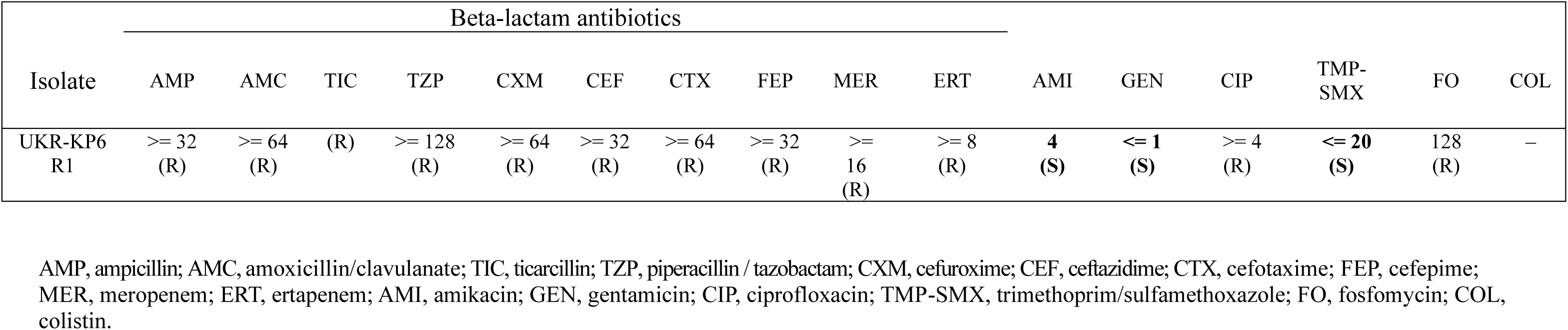
Antibiogram of UKR-KP6 resistant to phage Mavka.

Among several resistant isolates that retained the plasmid (4/9), multiple mutations were identified within the plasmid region annotated as the origin of replication (OriV), as well as a 189 bp insertion within a hypothetical gene (Supplementary Table 11). Because mutations in OriV can compromise plasmid replication and maintenance, these alterations may contribute to plasmid instability and explain the observed plasmid loss in a subset of isolates.

## Discussion

Recent surveillance studies have tracked multidrug-resistant organisms (MDROs) in countries receiving patients from Ukraine, but it remains challenging to determine whether these MDROs originated in Ukraine or were acquired during or after displacement to hospitals across Europe. We obtained problematic isolates belonging to the species *P. stuartii*, *P. aeruginosa*, *P. mirabilis*, *A. baumannii* and *K. pneumoniae* directly from the main military hospital in Kyiv in order to isolate and develop adequate therapeutic phages against them. Due to the predominance and initial assessment of resistance and virulence, we prioritized the comprehensive characterization of the *K. pneumoniae* isolates The initial *K. pneumoniae* isolates belonged to sequence types ST39, ST307, and ST395, with subsequent expansion to include ST101 and ST147 in 2025. Three isolates initially classified as ST23 (KL57) were reanalyzed using cgMLST and identified as belonging to clonal group CG10146, indicating they were more genetically distant from the typical ST23 lineage.

In February 2024, the European Centre for Disease Prevention and Control (ECDC) reported on a distinct *K. pneumoniae* lineage (ST23-KL57) which was first detected in Moscow, Russia in 2014^32^. Subsequent isolates were epidemiologically linked to Belarus, Russia, or Ukraine, with over half (6/11) of the recent isolates (2022-2023) primarily associated with Ukraine^32^. In our analysis of the available genomes in Pathogenwatch within the ST23-KL57 group, we found that 11 out of 28 isolates (excluding those described here) had origins in Ukraine. Among the remaining 17 isolates not originating from Ukraine, six were linked to Russia. These Russian isolates generally showed low resistance scores, high virulence scores, and earlier isolation dates. Notably, only one recent Russian isolate (SAMN18898210) exhibited both high virulence and resistance scores similar to Ukrainian isolates described in this paper. Based on cumulative data, including findings from a recent study on Ukrainian soldiers transferred to the burn wound center of the QAMH, this clonal group is increasingly prevalent in *K. pneumoniae* isolates among patients originating in Ukraine. The continued and worldwide spread of this clone could have severe implications, as recent isolates harbor the carbapenemases OXA-48 and NDM-1. The latter was identified within a large virulence plasmid alongside aerobactin, a key siderophore receptor and major virulence factor in *K. pneumoniae*, as well as the 16S rRNA methyltransferase ArmA, which confers high-level resistance to aminoglycosides.

Several *K. pneumoniae* isolates including those from the QAMH study and a German isolate grouped within ST395 which has been reported globally as an emerging threat, implicated in recent outbreaks in Turkey and Italy, and isolated from wounded patients from Ukraine^16,33–35^. The UKR isolates from this sequence type harbored a plasmid similar to one from an isolate taken from a patient in a hospital in Saint Petersburg, Russia, described in Starkova et al.^23^Here we report strains with this hybrid plasmid (phvKpST395_NDM1_1657) containing both NDM-1 and virulence genes in a K2 capsular serotype *K. pneumoniae* isolate in Ukraine.

The ST395 isolates described here carry NDM-1, ArmA, aerobactin and *rmpA2* within the same plasmid. In virulence testing the ST395 isolate UKR-KP6 was the most virulent in a *G. mellonella* model, acutely killing the larvae within 48 hours, similar to a classical hvKp strain. However, when tested in a mouse pneumonia model, this isolate was avirulent compared to a positive control in this model system. Of note, this positive control, which although has a lower virulence score of 4 (similar to the Ukrainian isolates) and does not carry the intact hypermucoviscous loci is still virulent in this pneumonia model. The Galleria model is useful for evaluating important virulence factors in *K. pneumoniae* particularly the role of capsule polysaccharide in pathogenesis^25,36^. Unlike the other ST395 isolates characterized here, UKR-KP6 has a predicted K2 capsule type, similar to the one described in St. Petersburg. This capsule type is strongly associated with virulence and hypervirulence in *K. pneumoniae*. Also note that another isolate from a Ukrainian soldier displaced in Germany (Figure 4; DE-KP27) was also determined to be ST395 KL2.

A separate QAMH surveillance study (Strachinaru et al., *unpublished*) identified several of these clonal groups in fecal and urine samples from displaced Ukrainian patients, highlighting their potential for carriage and dissemination, with success driven not only by AMR but also by enrichment of biosynthetic gene clusters encoding siderophores and RiPPs associated with virulence and interbacterial competition identified here. The continued global spread of these clonal groups, originating in Russia and Eastern Europe and now including 2025 isolates with acquired colistin resistance, underscores the urgent need for alternative therapeutic strategies.

Here we isolated diverse phages against all the UKR *K. pneumoniae* isolates. Several cases have demonstrated that phage therapy can be an effective treatment for resistant bacterial infections particularly when combined with antibiotics^6,7^. In one case, a 30-year-old victim of a suicide bombing in Brussels was successfully cured of *K. pneumoniae* wound infection using phages and antibiotics^37^. The success of this case was attributed to the synergistic interaction between phages and antibiotics both in biofilms and planktonic cells, and the use of pre-adapted phages tailored to the causative pathogen. Other cases have shown that phage-resistant bacterial strains, selected during phage therapy, may incur a fitness cost, leading to clinical improvement despite incomplete elimination of the pathogen^38^. Considering *K. pneumoniae* phages utilize receptors associated with antibiotic resistance (e.g., porins and efflux pumps) or virulence factors (e.g., siderophore receptors, LPS core, O-antigen, and capsule), phage resistance mutations could result in bacterial fitness costs or re-sensitization to antibiotics. This was demonstrated in a case where the patient’s strain developed resistance to three phages, accompanied by mutations in LPS and type IV pili, as well as in a critical component of the MexAB-OprM multidrug efflux pump and the DNA gyrase subunit A. The combination of these mutations leading to likely leading to re-sensitization to fluoroquinolones^6^.

In all phages assessed here, bacterial regrowth was observed in culture, although it was delayed. When we isolated phage-resistant bacteria, we identified mutations including large deletions in LPS and capsular biosynthesis genes. Notably, one isolate (UKR-KP6) had lost its hybrid plasmid, which resulted in re-sensitivity to aminoglycosides. Although the exact mechanism linking plasmid loss to phage resistance remains unclear, the capsule dependence of phage Mavka provides a plausible explanation. The hybrid plasmid carries *rmpA2*, a positive regulator of capsule synthesis, and its loss could prevent capsule upregulation or reversion to a more mucoid phenotype. Nonetheless, the resistor that lost the plasmid remained virulent in the *G. mellonella* model, suggesting that if the K2 capsule type is hypervirulent in this system, some level of capsule expression is still maintained.

As part of the Phage Sp UKR collaborative effort, Belgian defense is helping to develop phage therapy capabilities in a military hospital in Kyiv by providing phages, necessary equipment and training personnel to evaluate the usefulness of phages in the treatment of soldiers with difficult-to-treat infections. A complementary NATO STO report (Pirnay et al., unpublished) describes how phage could be deployed in contemporary military settings. Given the high specificity of *Klebsiella* phages, such applications would most likely be suited for Role/Echelon 3 or 4, where microbiological diagnostics are available. However, considering the predominance of highly clonal *K. pneumoniae* lineages identified in this study, it may also be feasible to develop phage preparations based on the available data for use at Role 1 and 2 levels, supporting patient care across the entire medical evacuation chain. These efforts have the potential to limit the continued spread of AMR, which is expected to persist beyond the end of the conflict.

## Methods

### Antibiotic susceptibility testing

Antibiotic susceptibility testing (AST) of *K. pneumoniae* isolates was performed using a VITEK 2 system (bioMéreux, Marcy L’Etoile, France) according to the manufacturer’s instructions. Categorization of minimum inhibitory concentrations (MICs) was based on the AST guidelines of the European Committee on Antimicrobial Susceptibility Testing (EUCAST).

### Bacterial growth

Isolates were grown in lysogeny broth (LB) from colonies plated on LB agar (1.5%). All bacteria were stored at -80°C in glycerol (15%) stocks. Class II biosafety cabinets were used under BSL2 conditions at all times when working with clinical isolates.

### Sequencing bacteria and phage

For the bacterial isolates genomic DNA was extracted using DNeasy UltraClean Microbial kit (Qiagen) using the protocol provided in the kit. Phage DNA was extracted using a protocol of pretreating DNase I and proteinase K to digest capsid proteins and release genomic DNA. This was concentrated using the Zymo DNA Clean & concentrator 5 kit. Genomic DNA was prepared for Nextera flex (Illumina) and sequenced on an Illumina MiniSeq. For the more recent isolates 2025 sequencing was performed on a NovaSeq 6000 system.

Nanopore sequencing (using rapid barcoding kit SQK-RBK114.24) was also performed on a MinION equipped with a R10.4.1 flow cell (Oxford Nanopore Technologies).

Basecalling was carried out with Guppy (v6.3.8, Oxford Nanopore Technologies). Sequencing of isolates LV-KP5546 and DE-KP427 was carried out using a PromethION 2 Solo (P2 Solo) device. Basecalling was performed using the Oxford Nanopore reads basecaller Dorado.

For the Illumina reads, quality of the reads was assessed using FastQC (v0.12.1, https://github.com/s-andrews/FastQC). Trimming was performed using Trimmomatic (Galaxy Version 0.39) for adapter clipping, quality trimming (LEADING:3 TRAILING:3 SLIDINGWINDOW:4:15) and minimum length exclusion (>50 bp)^39^. Porechop (Galaxy Version 0.2.4) was used for the nanopore reads. A hybrid assembly was constructed using Unicycler (Galaxy Version 0.5.). Unicycler generated assembly graphs which were visualized using Bandage software (v0.8.1). Annotation was performed using Bakta (web version)^40^.

Phage assembly was performed using Illumina reads only as described above. Annotation was performed using Pharokka (Galaxy Version 1.3.2).

All genomes are available in NCBI BioProject PRJNA1240947 for this study and for the QAMH burn wound center study in PRJNA1178460 (BE isolates).

### Genomic analysis

ABRicate (Galaxy Version 1.0.1; minimum DNA identity and coverage set to 90%) was used to determine resistant genes present in bacterial isolates^41^. The web application Pathogenwatch was used for the genomic assessment of *K. pneumoniae*^42^. Virulence scores (0-5), antibiotic resistance scores (0-3) and typing were determined using Kleborate (v2.2.0)^18^. The presence of virulence loci yersiniabactin/YbST, colibactin/CbST, and aerobactin/AbST,). was used to assign a virulence score: 0 = none present, 1 = yersiniabactin only, 2 = colibactin without aerobactin, 3 = aerobactin only, 4 = aerobactin and yersiniabactin without colibactin, and 5 = all three present. Kleborate screens for known AMR determinants using CARD AMR nucleotide database. Resistance scores were based on the presence of extended spectrum β-lactamases (ESBLs), carbapenemases, and colistin resistance: 0 = no ESBL or carbapenemase, 1 = ESBL without carbapenemase (regardless of colistin resistance), 2 = carbapenemase without colistin resistance (regardless of ESBL), and 3 = carbapenemase with colistin resistance (regardless of ESBL). K locus assignments were determined by Kaptive^43^. cgMLST schemes were provided by PubMLST, the Pasteur Institute, Enterobase and the cgMLST.org Nomenclature Server. The Pathogenwatch LIN code tool was used to infer *Klebsiella* lineage codes based on references from the Pasteur/PubMLST resource^44^. The sublineage and clonal group assignment were based on the third and fourth levels of the LIN code. Figure 1b was produced using Pathogenwatch and is a core distance-based neighbor-joining tree^45^. Figure 1a and Figure 1c were based on core genome assignment using Roary (Galaxy Version 3.13.0) (minimum identity for blastp 95%; percentage of isolates a gene must be to be in a core 99%)^46^. A maximum likelihood phylogenetic tree was created using FASTTREE (Galaxy Version 2.1.10) and visualized and annotated using iTOL (Version 6.9.1)^47^. All genomes used for Figure 1a are available on NCBI. All genomes used for the analysis are available on the Pathogenwatch website by sample accession # provided. Fig. 1c. Initial BGC detection and analysis of the eight isolates was performed with AntiSMASH software (v8.0.4)^48^ with detection strictness set into relaxed. BAGEL4^49^ and RiPP-Miner-Genome^50^ were employed for RiPP and bacteriocin detection.

Annotation of prototypic plasmids in ST23 and ST395 strains was manually curated using the Snapgene viewer tool, using for comparison the pLVPK virulence plasmid (Acc. No. AY378100) for UKR-KP3 and UKR-KP5 and the phvKpST395_NDM1_1657 plasmid (Acc. No. CP072809) for UKR-KP4, plasmids. PlasmidFinder was used to identify plasmid content (Supplementary Table 3). Phages were analyzed using the online tool VIRIDIC to determine genomic similarity^51^. A blastn search was performed to determine most similar phages. The lytic lifestyle of the phages was determined by a search using PhageAI platform and manual curation of the genome and determination of an absence of lysogeny related genes by searching for key terms like “integrase”, “excisionase”, “recombinase.”

### Virulence assays

The *Galleria mellonella* larvae infection model was performed as described with adjustments to dose^6^. Healthy larvae were weighed and separated into equal groups of 10 based on weight. Prior to this experiment optical density (OD) to CFU correlation calculation was performed to determine the CFU/mL based on OD. All cultures grew similarly. Overnight cultures were subcultured 1:100 and grown 37°C, shaking (250 rpm) until cultures reached OD_600_ of 0.25 – 0.55. Then cultures were adjusted to an OD_600_ of 0.1. The cultures were pelleted and resuspended in filter sterilized 1 X PBS buffer. The final dose was 1-5 X 10^4^ CFU. Larvae were infected in the hindmost left proleg with 10 µL of these suspensions. Following infection larvae were incubated at 37°C in the dark and monitored every 3 hours. Experiment for Figure 3 a,b was performed once. Experiment for Figure 3c, d was repeated twice.

Animal experiments were carried out in accordance with the National and European guidelines on animal experimentations. The project was evaluated by the ethics committee n°005 Charles Darwin (CETEA) and approved by the French Ministry of Higher Education and Research under the license number APAFIS #46235.

Eight-week-old female and male BALB/cJRj were purchased from Janvier Labs (Le Genest saint Isle, France). All mice were housed in the Biosafety Level 2+ animal facility and had free access to food and water and were under controlled light/dark cycle, temperature and humidity. Animals were handled with regard for alleviation of suffering. Animals were anesthetized using isoflurane and euthanized by cervical dislocation. Anesthetized mice were infected intranasally with 20 µL of either 10^6^, 10^7^ or 10^8^ UKR-KP6 or 10^7^ CG43 *K. pneumoniae* strains (N=6). Mice were weighted daily and their clinical score assessed twice daily. When the clinical score reached threshold values indicative of severe suffering, the animals were euthanized prior natural death consecutive of the infection. Mouse experiment was performed once with 3 males and repeated with 3 females. .

### Phage susceptibility testing

Susceptibility of bacterial strains towards the phages was tested using the spot test as described in Kutter et al.^52^. Freshly made suspensions of overnight agar cultures of the bacterial strains with adjusted OD_600_ of 0.2 (to an end concentration of 10^8^ CFU per mL) and in volume of 0.2 mL were added to 12 mL lukewarm (46°C) LB medium containing 0.6% agar (top agar) (and poured onto square (12 X 12 cm) Petri dishes (Greiner Bio-One, Vilvoorde, Belgium) containing media with 1.5% agar (bottom agar). LB Medium was purchased from Becton Dickinson (Erembodegem, Belgium) and agar from VWR International (Haasrode, Belgium). Droplets (10 μL) of serial dilutions of each of the considered phage solutions were spotted on the top agar layer. Petri dishes were incubated overnight at 32°C. The lysis zones produced by active phages in the bacterial lawn were examined. Next, for phages producing single plaques, the plaques were counted and the efficiency of plating (EOP) was determined as previously described^52^. The EOP for the considered bacterial strain was calculated by comparison of the phage titer on a propagating bacterial host and defined as the observed number of PFUs on the considered bacterial strain (as determined by spot test) divided by the number of PFUs on the reference bacterial host strain. In case the picture was unclear (for example, no obvious single plaques could be visualized) and the results were difficult to interpret, the double agar overlay method was used to determine the PFUs on the considered bacterial strains and on the bacterial reference host strains, to define EOP more precisely. For the double agar overlay method, one milliliter of phage lysate at appropriate dilutions (producing countable number of plaques 10^3^-10^1^ per mL) were added to a sterile 15 mL tube (Greiner Bio-One) and complemented with 0.1 mL of the considered bacterial suspension (to an end concentration of 10^8^ CFU per mL) and lukewarm (46°C) LB medium (purchased from Becton Dickinson) with 0.6% top agar (VWR International), to a total volume of 4 mL. This mixture was plated onto a round 90 mm diameter Petri dishes (Greiner Bio-One) filled with a bottom layer of LB and 1.5% agar (VWR International), and incubated overnight at 37°C. The plaques were examined and counted. The EOP was determined as previously described^53^. Phage antibiotic synergy assays were performed as described in Pirnay et al., 2023^6^.

### Phage isolation

Phages were isolated from sewage water of the Kyiv Military Hospital sampled in the period July – August 2023. Three milliliters of 10 X concentrated LB broth (Becton Dickinson), 1 mL of a suspension of 4 – 5 *K. pneumoniae* strains, at a concentration of 10^8^ CFU per mL in LB broth, and 26 mL of sewage water were mixed in a sterile 50 mL tube (Greiner Bio-One). The bacteria mixes consisted of 4 – 5 *K*.

*pneumoniae* strains isolated from Ukrainian soldiers in Kyiv (this study), Belgium (QAMH study), or Germany. This tube was incubated at 37°C for 3 – 4 hours. Six hundred microliters of chloroform (Sigma-Aldrich, Bornem, Belgium) were added, and the tube was slightly vortexed and then centrifuged for 20 min at 6,000 *g* and 4°C using a Sorvall Legend centrifuge (Thermo Fisher Scientific, Merelbeke, Belgium). The tube was further incubated for 1 hour at 4°C. Five milliliters of the supernatant was aspirated using a sterile 5 mL syringe (BD Plastipak, Becton Dickinson) and passed through a 0.45 µm membrane filter (Minisart, Sartorius, Vilvoorde, Belgium). The tubes were screened for the presence of plaques using the spot test, as described above. Tubes that showed plaque formation were further analyzed using the double agar overlay method, as described above. All plaques exhibiting different morphologies were touched with a sterile pipette tip, inoculated into 2 mL of sterile LB broth in 15 mL sterile tubes and incubated for 2 hours at 37°C. Then, 100 µL of chloroform were added and the tubes (one tube per plaque) were incubated for 1 hour at 4°C. For each tube, a dilution series (1:1 – 1:10^6^) was made in sterile 15 mL tubes filled with 0.9% NaCl solution. Each dilution was titrated using the agar overlay method (see above). Plates showing 1 – 10 plaques were further analyzed. Again, all plaques with different morphologies were touched with a sterile pipette tip, inoculated into 2 mL of sterile LB broth in 15 mL sterile tubes and incubated for 2 hours at 37°C. This cycle was repeated until one plaque morphotype was obtained (at least 5 times).

### Killing curve assay

Single colonies from freshly plated cultures of *K. pneumoniae* were grown in LB broth for four hours at 37°C, shaking (250 rpm). OD of the cultures was adjusted to 0.2 and seeded onto a 96 well plate (Greiner-flat bottom) so that 10^8^ CFU per well was added. Phages were added at a multiplicity of infection (MOI) of 1 (10^8^ PFU/well). Optical density was measured using a CLARIOstar® Plus Microplate Reader (BMG Labtech, Ortenberg, Germany) every 5 minutes under 300 rpm shaking conditions at 37°C for 24 hours.

Readings were blank (LB broth) adjusted. Phage killing curve was generating based on the average of 3 biological replicates.

Phage resistors were isolated by repeating the experiment above. Cultures were plated from each of the biological replicates. Single colonies were isolated from each of the biological replicates. The single colonies were restreaked three times then tested for resistance using a double agar-overlay assay as described above. Confirmed resistance (by absence of plaquing of that phage) was determined for 3 colonies of UKR-KP2 (R1-R3), two for UKR-KP3 (R1, R2), three for UKR-KP5.1 (R1-R3), nine for UKR-KP6 (R1-R9), 3 for UKR-KP7 (R1-R3) and two for UKR-KP8 (R1, R2).

### Plasmid stability test

Stability tests were performed as described^54^ with some modifications. Cultures were incubated at 37°C shaking (250 rpm) in LB for 24 hours, then serially diluted and plated in antibiotic-free LB agar and LB agar with gentamicin (2 μg/mL). Retention rate was calculated by dividing number of colonies that grew on gentamicin by number of colonies that grew on LB without antibiotic. In order to confirm 10 colonies were PCR verified using primers for the *armA* gene. This was performed using two biological replicates.

### Statistics

The log-rank test with Bonferroni correction for multiple comparisons (GraphPad Version 10.4.1) was used for *G. mellonella* survival curve comparisons.

## Acknowledgements

We thank the Institute Pasteur teams for the curation and maintenance of BIGSdb-Pasteur databases at http://bigsdb.pasteur.fr/. Thanks to Dr. Franklin Nobrega for advice and assistance with a research question.

CC, GS, and MM are supported by Grant HFM 24-17 of the Royal Higher Institute for Defence. SG, MN, RT, VL, CE, JW and RL are supported within the framework of the JPIAMR - Joint Programming Initiative on Antimicrobial Resistance by the KLEOPATRA project. MN gratefully thanks the Higher Education Commission of Pakistan for her scholarship no: PD/HEC/HRD/OSS-III/BIg-B2/2022/19801.

## Contributions

S.G. and M.N. performed experiments, analyzed data, generated figures, wrote and edited the manuscript. A.K., O.R., O.S., and V.T. provided samples and advice for the manuscript. A. C. generated Fig. 2 and provided analysis of plasmid content of the strains and edited the manuscript. P.B. provided strains and edited the manuscript. R.T., V.L. and C.E. performed mouse experiments (Figure 3), analyzed the data, edited the manuscript. E.D. contributed experimentally to Figure 5. D.S. provided strains and edited the manuscript. P.J. provided strains and edited the manuscript. G.S., C.C., J.G., and M.M. isolated phages and contributed experimentally to Figure 5 and Supplementary Figure 2. M.M. edited the manuscript.

A.S. performed MIC assays for Tables 1-3. K.R. and V.A. provided samples and edited the manuscript. J.W. reviewed and edited the manuscript. T.L. assisted in performing sequencing for Figure 4. E.V. provided analysis of secondary metabolites for Supplementary Table 4. R.L. provided guidance and resources throughout the project and reviewed and edited the manuscript. J.P. provided guidance and resources, led the project and reviewed and edited the manuscript. All authors reviewed the manuscript.

