## Supplementary Tables for "Genomic epidemiology of conflict-associated *Klebsiella pneumoniae* in Ukraine reveals a high-risk clone with a hybrid plasmid vulnerable to phages"

**Supplementary Table 1** Resistance genes present in UKR isolates.

| Isolate | K Locus | Antibiotic types | | | | | | |
|---|---|---|---|---|---|---|---|---|
| | | Aminoglycosides | Quinolones | Tetracyclines | Macrolides | Sulfonamides | Antifolates | Others[1.] |
| UKR-KP2 | KL23 | *ant(2'')-Ia,aph(3'')-Ib, aph(6)-Id, aac(6')-Ib, armA, rmtC* | *qnrS1, aac(6')-Ib-cr ,* | *tet(A)* | *msr(E), mph(E)* | *sul1, sul2* | | *arr-3, catA1, cmlA1* |
| UKR-KP3 | KL57 | *aph(3')-VI, armA* | *qnrS1* | | *msr(E), mph(E), mph(A)* | *sul1, sul2* | *dfrA5* | |
| UKR-KP4 | KL108 | *aac(3)-IIa, aph(3')-VI, armA* | *qnrS1, aac(6')-Ib-cr* | *tet(A)* | *msr(E), mph(E), mph(A)* | *sul1, sul2* | *dfrA5, dfrA1* | *catA1* |
| UKR-KP5.1 | KL57 | *aph(3')-VI, armA* | *qnrS1* | | *mph(A), msr(E), mph(E)* | *sul1, sul2* | *dfrA5* | |
| UKR-KP5.2 | KL57 | *aph(3')-VI, armA* | *qnrS1* | | *mph(A), msr(E), mph(E)* | *sul1, sul2* | *dfrA5* | |
| UKR-KP6 | KL2 | *aph(3')-VI, armA* | *qnrS1, aac(6')-Ib-cr* | | *mph(E), msr(E), mph(A)* | *sul1, sul2* | *dfrA5* | *catA1* |
| UKR-KP7 | KL23 | *ant(2'')-Ia, ant(3'')-Ia, aac(6')-Ib aph(6)-Id, aph(3'')-Ib, armA, rmtC* | *qnrS1, aac(6')-Ib-cr* | *tet(A)* | *mph(E), msr(E),* | *sul1, sul2* | | *catA1, cmlA1, arr-3* |
| UKR-KP8 | KL102 | *aph(3'')-Ib, aph(6)-Id, aac(6')-Ib* | *aac(6')-Ib-cr, qnrB1* | *tet(A)* | | *sul2* | *dfrA14* | |

[1.] All strains carried the *oqxA*, *oqxB* and *fosA* genes intrinsic in *K. pneumoniae*.

**Supplementary Table 2**

Bela-lactamase genes present in UKR isolates.

| Isolates | *bla*$_{KPC-2}$ | *bla*$_{NDM-1}$ | *bla*$_{TEM-1B}$ | *bla*$_{CMY-16}$ | *bla*$_{CTX-M15}$ | *bla*$_{CTX-M55}$ | *bla*$_{OXA-1}$ | *bla*$_{OXA-10}$ | *bla*$_{OXA-48}$ |
|---|---|---|---|---|---|---|---|---|---|
| UKR-KP2 | ■ | ■ | ■ | ■ | ■ | | ■ | ■ | |
| UKR-KP3 | | ■ | | | | ■ | | | ■ |
| UKR-KP4 | | ■ | ■ | | ■ | | ■ | | |
| UKR-KP5.1 | | ■ | | | | ■ | | | ■ |
| UKR-KP5.2 | | ■ | | | | ■ | | | ■ |
| UKR-KP6 | | ■ | ■ | | ■ | | ■ | | ■ |
| UKR-KP7 | ■ | ■ | ■ | ■ | ■ | | ■ | ■ | |
| UKR-KP8 | | | ■ | | ■ | | ■ | | ■ |

**Supplementary Table 3**

Virulence and antibiotic resistance score of UKR isolates.

| Isolates | Virulence | Antibiotic resistance |
|---|---|---|
| UKR-KP2 | 4 | 3 |
| UKR-KP3 | 4 | 3 |
| UKR-KP4 | 3 | 3 |
| UKR-KP5.1 | 4 | 3 |
| UKR-KP5.2 | 4 | 3 |
| UKR-KP6 | 4 | 3 |
| UKR-KP7 | 4 | 3 |
| UKR-KP8 | 1 | 3 |

**Supplementary Table 4** Secondary metabolite clusters identified with AntiSMASH

| | Type/Class | Location | Most similar known cluster |
|---|---|---|---|
| 1 | T1PKS, NRP-metallophore, NRPS | UKR-KP2: 1,801,828-1,865,027; UKR-KP3: 1,702,339-1,765,538; UKR-KP5.1: 1,831,830-1,895,029; UKR-KP5.2: 1,824,830-1,888,029; UKR-KP6: 2,000,702-2,063,901; UKR-KP7: 442,859-506,058; UKR-KP8: 1,808,745-1,871,944 | Yersiniabactin |
| 2 | Redox-cofactor | UKR-KP2: 2,564,900-2,587,065; UKR-KP3: 2,372,704-2,394,869; UKR-KP4: 2,626,299-2,648,464; UKR-KP5.1: 2,502,195-2,524,360; UKR-KP5.2: 2,495,195-2,517,360; UKR-KP6: 2,703,330-2,725,495; UKR-KP7: 780,152-802,317; UKR-KP8: 2,506,300-2,528,465 | - |
| 3 | Azole-containing-RiPP | UKR-KP2: 3,475,785-3,502,054; UKR-KP3: 3,284,342-3,310,611; UKR-KP4: 1,714,876-1,741,144; UKR-KP5.1: 3,412,681-3,438,950; UKR-KP5.2: 3,407,349-3,433,618; UKR-KP6: 3,587,987-3,614,255; UKR-KP7: 1,691,038-1,717,307; UKR-KP8: 3,372,049-3,398,318 | - |
| 4 | NRP-metallophore, NRPS | UKR-KP2: 3,810,323-3,864,095; UKR-KP3: 3,619,941-3,673,967; UKR-KP4: 1,322,512-1,376,420; UKR-KP5.1: 3,748,280-3,802,306; UKR-KP5.2: 3,742,948-3,796,974 UKR-KP6: 3,952,822-4,006,730; UKR-KP7: 1,691,038-1,717,307; UKR-KP8: 3,718,188-3,772,330 | Enterobactin |
| 5 | Terpene precursor | UKR-KP2: 4,147,608-4,168,507; UKR-KP3: 3,838,961-3,859,860; UKR-KP4: 1,007,427-1,028,326; UKR-KP5.1: 3,967,300-3,988,199; UKR-KP5.2: 3,961,968-3,982,867 UKR-KP6: 4,298,733-4,319,632; UKR-KP7: 2,362,753-2,383,652; UKR-KP8: 4,005,769-4,026,668 | - |
| 6 | RiPP-like | UKR-KP2: 5,322,349-5,332,972; UKR-KP3: 4,983,038-4,993,661; UKR-KP4: 5,340,802-5,351,425; UKR-KP5.1: 5,111,371-5,121,994; UKR-KP5.2: 5,106,043-5,116,666 UKR-KP6: 5,515,145-5,525,768; UKR-KP7: 2,361,224-2,371,847; UKR-KP8: 5,255,020-5,265,643 | - |
| 7 | NI-siderophore | UKR-KP2: 249,894-282,356; UKR-KP3: 127,654-160,116 UKR-KP4: 122,070-154,532; UKR-KP5.1: 98,759-131,221 UKR-KP5.2: 98,759-131,221; UKR-KP6: 58,796-91,258 UKR-KP7: 242,568-275,030 | Aerobactin |
| 8 | RiPP-like | UKR-KP2: 1-9,294; UKR-KP3: 1-8,306; UKR-KP5.1: 1,865-9,294; UKR-KP5.2: 1-6,387; UKR-KP7: 1-9,294 UKR-KP8: 2,161-13,846 | - |

**Supplementary Table 5** Plasmid content of UKR isolates.

| ST | Isolate | Plasmids | Size (bp) | Contig | | Plasmids | Size (bp) | Contig |
|---|---|---|---|---|---|---|---|---|
| ST39 | UKR-KP2 | IncHI1B(pNDM-MAR)_1__JN420336 | 325067 | 2 | UKR-KP7 | IncHI1B(pNDM-MAR)_1__JN420336 | 317740 | 3 |
| | | IncC_1__JN157804 | 152850 | 3 | | IncC_1__JN157804 | 152850 | 4 |
| | | IncFIB(pKPHS1)_1__CP003223 | 110027 | 4 | | IncFIB(pKPHS1)_1__CP003223 | 110027 | 5 |
| | | IncFII(K)_1__CP000648 | 102252 | 5 | | IncFIB(pQil)_1__JN233705, IncFII(K)_1__CP000648 | 102247 | 6 |
| | | ColRNAI_1__DQ298019 | 9294 | 6 | | ColRNAI_1__DQ298019 | 9294 | 7 |
| | | Col_pB8919_5_CP094996.1 | 4352 | 7 | | Col_pB8919_5_CP094996.1 | 4352 | 8 |
| ST23 | UKR-KP3 | repB_KLEB_VIR_AP006726, RepB_pC39_CP061702.1 | 225640 | 2 | UKR5.1/5.2 | repB_KLEB_VIR_AP006726, RepB_pC39_CP061702.1 | 243065 | 2 |
| | | IncFIA(HI1)_1__AF250878, IncFII(K)_1__CP000648 | 88127 | 3 | | IncFIA(HI1)_1__AF250878, IncFII(K)_1__CP000648 | 88141 | 3 |
| | | IncFIIhvKP319_3_CP101767.1 | 83831 | 4 | | IncFIIhvKP319_3_CP101767.1 | 83831 | 4 |
| | | IncFII_1__AY458016 | 44324 | 6 | | IncFII_1__AY458016 | 48972 | 5 |
| | | ColRNAI_1__DQ298019 | 9294 | 7 | | ColRNAI_1__DQ298019 | 9294 | 6 |
| | | Col(pHAD28)_1__KU674895 | 4429 | 8 | | Col(pHAD28)_2__KU674895 | 4429 | 7 |
| | | Col(pHAD28)_2__KU674895 | 4167 | 9 | | Col(pHAD28)_1__KU674895 | 4167 | 8 |
| | | ColpVC_1__JX133088 | 1934 | 12 | | ColpVC_1__JX133088 | 1934 | 9 |
| ST395 | UKR-KP4 | IncHI1B(pNDM-MAR)_1__JN420336, IncFIB(pNDM-Mar)_1__JN420336 | 346894 | 2 | UKR-KP6 | IncHI1B(pNDM-MAR)_1__JN420336, IncFIB(pNDM-Mar)_1__JN420336 | 356463 | 2 |
| | | IncR_1__DQ449578, IncN_1__AY046276 | 67290 | 3 | | IncFIB(K)_1__JN233704, IncFII(K)_1__CP000648 | 224939 | 3 |
| | | Col(pHAD28) | 3511 | 4 | | IncL_1__JN626286 | 63589 | 4 |
| | | | | | | Col_CriePir75_CP063021.1 | 5010 | 5 |
| ST307 | UKR-KP8 | IncFIB(K)_1__JN233704 | 152998 | 2 | | | | |
| | | IncL_1__JN626286 | 54230 | 3 | | | | |
| | | ColRNAI_1__DQ298019 | 14840 | 4 | | | | |

**Supplementary Table 6** Resistance genes present in 2025 UKR isolates.

| Isolate | Antibiotic types | | | | | | |
|---|---|---|---|---|---|---|---|
| | Aminoglycosides | Quinolones | Tetracyclines | Macrolides | Sulfonamides | Antifolates | Others[1.] |
| UKR-KP136 | *aph(3')-VI, armA, aac(3)-IIa* | *qnrS1* | | *mph(A), mph(E), msr(E)* | *sul1, sul2* | *dfrA5* | |
| UKR-KP219 | *aph(3')-VI, armA, aac(3)-IIa* | *qnrS1* | | *mph(A), mph(E), msr(E)* | *sul1, sul2* | *dfrA5* | |
| UKR-KP234 | *aph(3')-VI, armA, aac(3)-IIa* | *qnrS1* | | *mph(A), mph(E), msr(E)* | *sul1, sul2* | *dfrA5* | |
| UKR-KP328 | *aph(3')-VI, ant(2'')-Ia, ant(3'')-Ia, aac(6')-Ib* | *qnrS1, aac(6')-Ib-cr* | | | *sul1* | | *catA1, catB3, ARR-3* |
| UKR-KP335 | *aph(3')-VI, ant(2'')-Ia, ant(3'')-Ia, aac(6')-Ib* | *qnrS1, aac(6')-Ib-cr* | | | *sul1* | | *catA1, catB3, ARR-3* |
| UKR-KP336 | *aph(3')-VI, armA, aac(3)-IIa* | *qnrS1* | | *mph(A), mph(E), msr(E)* | *sul1, sul2* | *dfrA5* | |
| UKR-KP370 | *aph(3')-VI, armA, aac(3)-IIa* | *qnrS1* | | *mph(A), mph(E), msr(E)* | *sul1, sul2* | *dfrA5* | |
| UKR-KP371 | *aph(3')-VI, ant(3'')-Ia, aac(6')-Ib* | *qnrS1,* | | *mph(A), mph(E), msr(E)* | *sul1, sul2* | *dfrA5* | |
| UKR-KP386 | *ant(2'')-Ia, ant(3'')-Ia, aac(3)-IIa, aph(3'')-Ib, aph(6)-Id* | *qnrB1, aac(6')-Ib-cr* | *tet(A)* | | *sul1, sul2* | *dfrA14* | *catA1* |
| UKR-KP627c | *aph(3')-VI, armA, aac(3)-IIa* | *qnrS1* | | *mph(A), mph(E), msr(E)* | *sul1, sul2* | *dfrA5* | |
| UKR-KP636 | *aph(3')-VI, ant(2'')-Ia, ant(3'')-Ia, aac(6')-Ib* | *qnrS1, aac(6')-Ib-cr* | | | *sul1* | | *catA1, catB3, ARR-3* |
| UKR-KP697 | *ant(2'')-Ia, ant(3'')-Ia* | *qnrS1, aac(6')-Ib-cr* | | | *sul1* | | *catA1* |
| UKR-KP1012 | *aph(3')-VI, armA, aac(3)-IIa* | *qnrS1* | | *mph(A), mph(E), msr(E)* | *sul1, sul2* | *dfrA5* | |
| UKR-KP1131 | *armA* | *qnrS1* | | *mph(A), mph(E), msr(E)* | *sul2* | | |
| UKR-KP1132 | *aph(3')-VI, armA, aac(3)-IIa* | *qnrS1* | | *mph(A), mph(E), msr(E)* | *sul1, sul2* | *dfrA5* | |
| UKR-KP1720 | *rmtF, armA, ant(2'')-Ia, ant(3'')-Ia* | | | *mph(E), msr(E)* | *sul1* | | *catA1, ARR-3* |

1. All strains carried the *oqxA*, *oqxB* and *fosA* genes intrinsic in *K. pneumoniae*.

**Supplementary Table 7** Plasmid content of 2025 UKR isolates.

| ST | Isolate | Plasmids | Size (bp) | Contig | | Plasmids | Size (bp) | Contig |
|---|---|---|---|---|---|---|---|---|
| ST39 | UKR-KP697 | IncHI1B(pNDM-MAR)__JN420336 | 314381 | 2 | | | | |
| | | IncFII(K)_1__CP000648 | 102252 | 4 | | | | |
| | | IncFIB(pQil)__ JN233705 | | | | | | |
| | | ColRNAI_1__DQ298019 | 9294 | 5 | | | | |
| | | Col_pB8919_5_CP094996.1 | 4352 | 6 | | | | |
| ST23 | UKR-KP136 | repB_KLEB_VIR_AP006726, RepB_pC39_CP061702.1 | 225757 | 2 | UKR-KP219 | repB_KLEB_VIR_AP006726, RepB_pC39_CP061702.1 | 259768 | 6 |
| | | IncFIA(HI1)_1__AF250878, IncFII(K)_1__CP000648 | 88154 | 3 | | IncFIA(HI1)_1__AF250878, IncFII(K)_1__CP000648 | 88153 | 9 |
| | | IncFII(pSDP9R)__MW940621 | 83831 | 4 | | IncFII(pSDP9R)__MW940621 | 58434 | 5 |
| | | IncFII_1__AY458016 | 47744 | 5 | | IncFII_1__AY458016 | 47743 | 4 |
| | UKR-KP234 | repB_KLEB_VIR_AP006726, RepB_pC39_CP061702.1 | 243191 | 2 | UKR-KP336 | repB_KLEB_VIR_AP006726, RepB_pC39_CP061702.1 | 225757 | 2 |
| | | IncFIA(HI1)_1__AF250878, IncFII(K)_1__CP000648 | 88154 | 3 | | IncFIA(HI1)_1__AF250878, IncFII(K)_1__CP000648 | 88154 | 3 |
| | | IncFII(pSDP9R)__MW940621 | 83831 | 4 | | IncFII(pSDP9R)__MW940621 | 83831 | 4 |
| | | IncFII_1__AY458016 | 47744 | 5 | | IncFII_1__AY458016 | 47744 | 5 |
| | UKR-KP370 | repB_KLEB_VIR_AP006726, RepB_pC39_CP061702.1 | 243191 | 2 | UKR-KP627c | repB_KLEB_VIR_AP006726, RepB_pC39_CP061702.1 | 225757 | 2 |
| | | IncFIA(HI1)_1__AF250878, IncFII(K)_1__CP000648 | 88154 | 3 | | IncFIA(HI1)_1__AF250878, IncFII(K)_1__CP000648 | 88154 | 3 |
| | | IncFII(pSDP9R)__MW940621 | 83831 | 4 | | IncFII(pSDP9R)__MW940621 | 83831 | 4 |
| | | IncFII_1__AY458016 | 47762 | 5 | | IncFII_1__AY458016 | 47744 | 5 |
| | UKR-KP1131 | repB_KLEB_VIR_AP006726, RepB_pC39_CP061702.1 | 119570 | 2 | UKR-KP1132 | repB_KLEB_VIR_AP006726, RepB_pC39_CP061702.1 | 225749 | 2 |
| | | IncFIA(HI1)_1__AF250878, IncFII(K)_1__CP000648 | 88154 | 3 | | IncFIA(HI1)_1__AF250878, IncFII(K)_1__CP000648 | 88154 | 3 |
| | | IncFII(pSDP9R)__MW940621 | 83831 | 4 | | IncFII(pSDP9R)__MW940621 | 83831 | 4 |
| | | IncFII_1__AY458016 | 44324 | 5 | | IncFII_1__AY458016 | 47762 | 5 |
| | | ColRNAI__DQ298019 | 9294 | 6 | | | | |
| ST307 | UKR-KP386 | IncHI1B(pNDM-MAR)__JN420336 | 295484 | 2 | | | | |
| | | IncFIB(pNDM-Mar) | | | | | | |
| | | IncFIB(K)__JN233704 | 158904 | 3 | | | | |
| ST147 | UKR-KP328 | IncHI1B(pNDM-MAR)__JN420336 | 299622 | 3 | UKR-KP335 | IncHI1B(pNDM-MAR)__JN420336 | 264446 | 2 |
| | | IncFIB(pQil)__JN233705 | 54064 | 9 | | IncFIB(pQil)__JN233705 | 93664 | 4 |
| | | | | | | IncR_DQ449578 | | |
| | | IncR_DQ449578 | 39748 | 10 | | IncFII(K)__CP000648 | 86237 | 5 |
| | | | | | | IncFIA(HI1)__AF250878 | | |
| | UKR-KP371 | IncHI1B(pNDM-MAR)__JN420336 | 349357 | 2 | UKR-KP636 | IncHI1B(pNDM-MAR)__JN420336 | 264447 | 4 |
| | | IncR_DQ449578 | 51593 | 4 | | IncFII(K)__CP000648 | 86237 | 9 |
| | | | | | | IncFIA(HI1) __AF250878 | | |
| | | ColpVC _JX133088 | 2058 | 6 | | IncFIB(pQil)__JN233705 | 54064 | 11 |
| | | | | | | IncR_DQ449578 | 39600 | 12 |
| ST101 | UKR-KP1720 | IncHI1B(pNDM-MAR) _JN420336 | 310912 | 2 | | | | |
| | | IncFIB(pNDM-Mar)__JN420336 | | | | | | |
| | | IncFIB(K)__JN233704 | 186859 | 3 | | | | |
| | | IncFIA(HI1)__AF250878 | 72419 | 4 | | | | |
| | | IncR_DQ449578 | | | | | | |
| | | ColRNAI__DQ298019 | 9294 | 5 | | | | |
| | | Col440II__CP023921 | 5359 | | | | | |

**Supplementary Table 8** Phage Blastn results to other phages.

| Phage | Blastn hit | Accession # | Identity (%) | Query cover (%) |
|---|---|---|---|---|
| Ukrainka | KpV17Sklif | PQ663821 | 93.8 | 89 |
| Mavka | vB_KpnP_NER40 | MZ602146 | 90.18 | 89 |
| Odessa | RCIP0041 | OR532835 | 92.7 | 70 |
| Lesya | vB_KpP_FBKp27 | NC_068555 | 95 | 87 |
| Dzikanka | Sweeny | NC_049839 | 94.16 | 92 |

**Supplementary Table 9**

Single nucleotide polymorphisms (SNPs), small insertions and deletions (INDELS) identified in phage-resistant mutants.

| Isolate | Phage | Mutation | Gene affected | Frequency |
|---|---|---|---|---|
| UKR-KP2 | Ukrainka | stop_gained c.793G>T p.Glu265* | *arnC* | 1/2 |
| UKR-KP3 | Mikola | frameshift_variant c.1102_1105delGGCA p.Gly368fs | *lamB* | 2/2 |
| UKR-KP6 | Mavka | missense_variant c.1600G>T p.Gly534Cys | *wzC* | 1/9 |
| | | missense_variant c.1200G>T p.Met400Ile | *wcaJ* | 1/9 |

**Supplementary Table 10**

Large deletions identified in phage-resistant mutants.

| Isolate | Phage | Size of deletion | Gene(s) affected | Frequency |
|---|---|---|---|---|
| UKR-KP6 | Mavka | 356,463 bp | *iuc*, *armA*, $_{bla}$*NDM-1*, *msrE*, *mphA*, *mphE*, *qnrS1*, *Sul1*, *Sul2*, *dfrA5* | 3/9 |
| UKR-KP7 | Ukrainka | 9,273 bp | *ugd*, *rfbC*, *rfbD*, *rffH*, *rffG*, *gnd* | 1/2 |
| | | 5,741-5,755 bp | *wzC*, wz*A*, wz*I* | 2/2 |

**Supplementary Table 11**

| Bacteria | Mutation | Region affected | Annotation | Frequency |
|---|---|---|---|---|
| KP6 | Multiple SNPs | Contig 2 256802..257850 | Origin of replication | 4/9 |
| KP6 | 189 bp insertion | Contig 2 268493 | VWA domain containing protein | 4/9 |
