## Supplementary Figures for "Genomic epidemiology of conflict-associated *Klebsiella pneumoniae* in Ukraine reveals a high-risk clone with a hybrid plasmid vulnerable to phages"

### Supplementary Figure 1 VIRIDIC (Virus Intergenomic Distance Calculator)

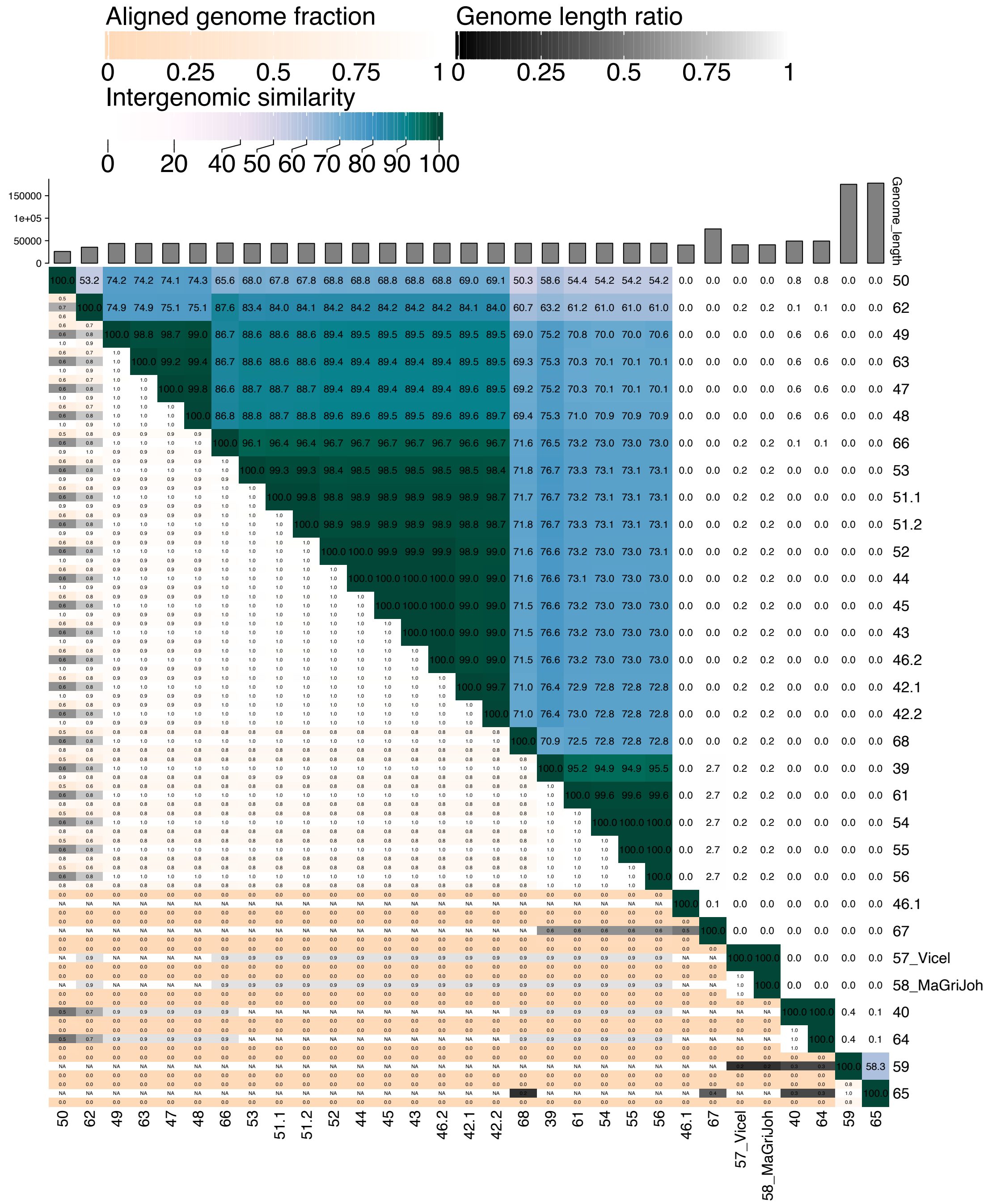

### Supplementary Figure 2

**a**

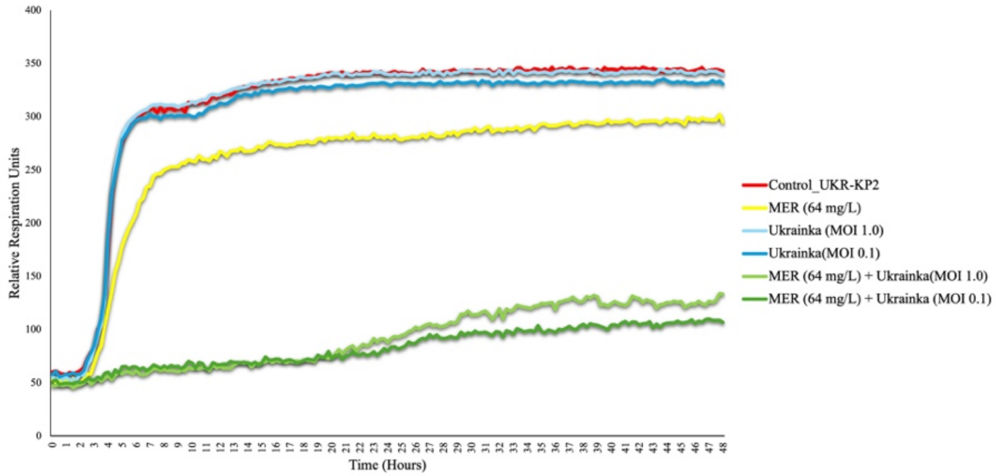


**b**

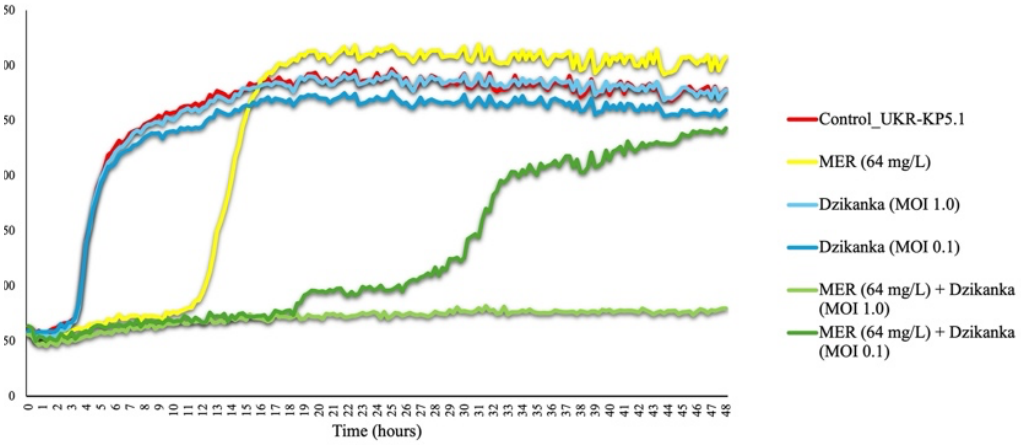


**c**

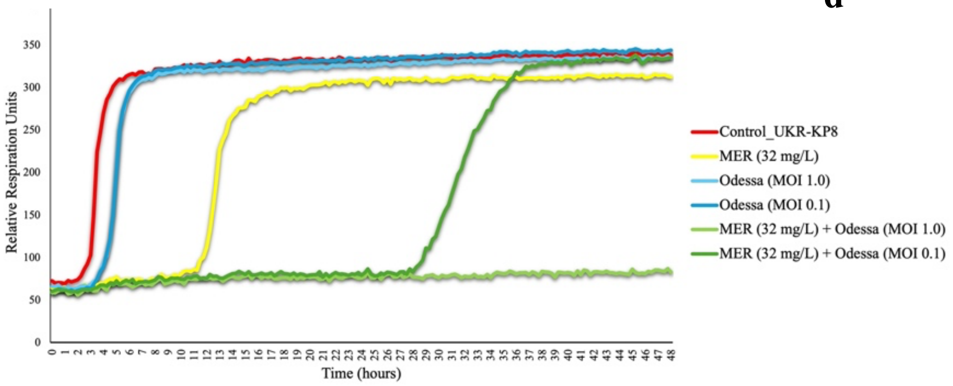


**d**

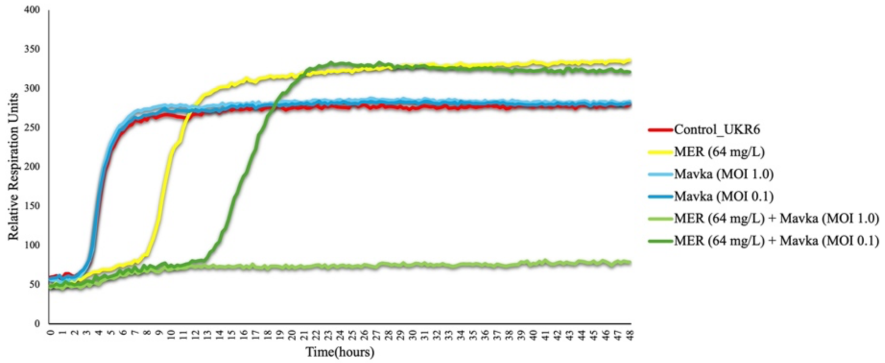

**Supplementary Figure 2 Synergistic interaction of UKR phages with meropenem** Omnilog data of bacterial growth for 48 hours using the antibiotic meropenem with and without phages a. Ukrainka (UKR-KP2), b. Dzikanka (UKR-KP5.1), c. Mavka (UKR-KP6) and d. Odessa (UKR-KP8). The median is represented of 3 replicates (N=3).

**Supplementary Figure 3**

**a**

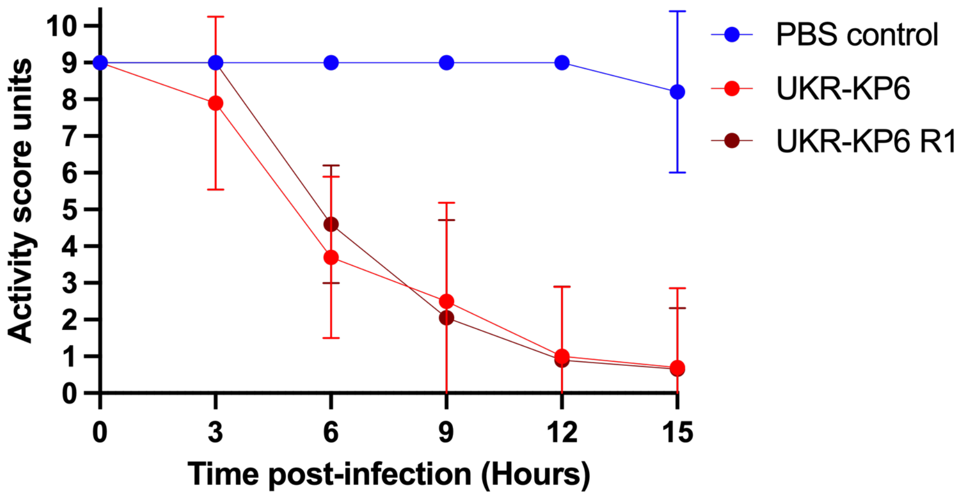


**b**

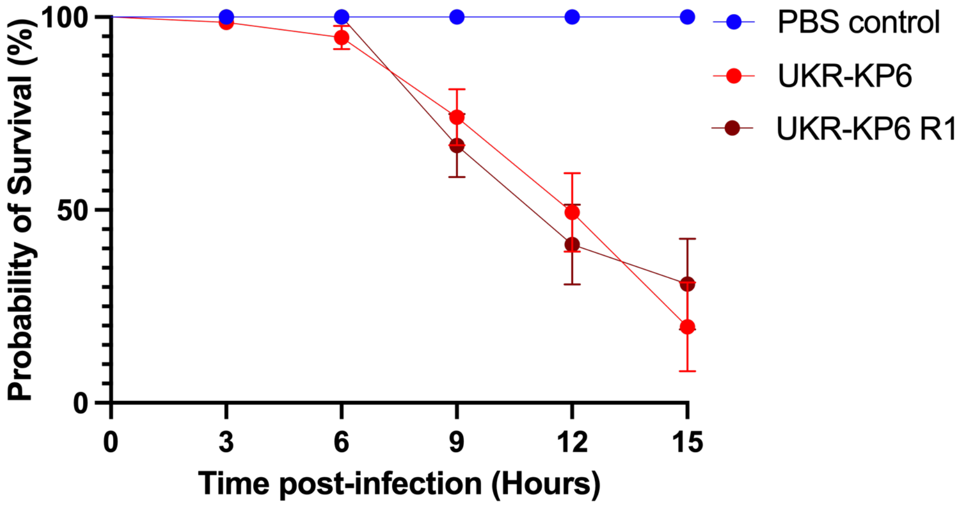


**Supplementary Figure 3 Virulence assessment of UKR-KP6 and phage-resistant mutant UKR-KP6 R1. a,** *G. mellonella* larvae (n = 10) were injected with *K. pneumoniae* isolates at a dose of $10^4$ CFU or with PBS and incubated at a temperature of 37°C. The larvae were monitored every 3 hours to assess for activity. **b,** Probability of survival. Mean and standard deviation are represented. This experiment was performed using 2 biological replicates.
